# A High-Performance and Recoverable DNA Origami Rotary Motor

**DOI:** 10.1101/2025.04.04.645735

**Authors:** Haggai Shapira, Breveruos Sheheade, Samrat Basak, Dinesh Khara, Eyal Nir

## Abstract

Inspired by biological molecular machines, we developed a highly processive DNA origami rotary motor. The rotor consists of two disk-shaped DNA origamis that are connected by a single-stranded swivel that allows free rotation but prevents rotor dissociation in the event of operational error. The rotor is propelled by two bipedal walkers that stride on a circular DNA track propelled by a previously optimized DNA propulsion mechanism called ‘fuel before antifuel’. The DNA fuel and antifuel strands are delivered by a microfluidic device. The rotation is monitored by a single-molecule, light scattering, defocused imaging technique; light scattered from a gold nanorod attached to the rotor enables high angular and temporal resolution analyses of rotor orientation. Imaging movies show individual rotors performing 96 individual steps, corresponding to 8 full rotor revolutions, with direction determined by the fuel and antifuel sequence. In cases of operational errors, which result in free Brownian rotation, the rotors were able to recover and continue rotating as commanded. Our origami-based rotary motor design, microfluidics-based control system, and high-resolution monitoring of rotor orientation will facilitate the development of DNA-based machines driven by autonomous propulsion.

## Introduction

Biological molecular motors made of proteins, which function in many biological processes, often operate with remarkably high processivity and speed. For example, a kinesin bipedal walker performs hundreds of steps within seconds before dissociating from the microtubule track^1^, and the F1-ATPase and bacterial flagellum rotary motors can rotate almost indefinitely^2, 3, 4, 5^. These biological rotary motors are encapsulated within a membrane that prevents rotor dissociation in the event of operational errors and allows them to fully recover.

Inspired by biological motors, numerous artificial molecular motors have been developed^6, 7^. Specifically, the DNA nanotechnology and DNA origami techniques^8, 9, 10, 11, 12, 13, 14, 15^ have allowed rational development of structurally complex, programmable and functional molecular machines^16, 17, 18, 19, 20, 21, 22^, and, in recent years, these methods have been used to manufacture several new types of rotary devices^23, 24, 25, 26, 27, 28, 29^. These rotary devices employ various means of actuation including electric and magnetic fields^23, 24, 28^, biological molecular machines^25^, DNA-zymes^26^, and DNA fuels^26^. Impressively, a DNA “windmill” based on ion flow performed up to 200 revolutions before failure^29^. However, a highly processive DNA origami rotary motor powered by DNA fuels that performs more than few steps has yet to be demonstrated.

In contrast to electric and magnetic fields, light, ion flow, and other more specific powering methods, the richness of DNA sequences and their orthogonality allows the development of many different powering mechanisms, including ones that operate in parallel, and even allow the development of autonomously operating machines as was demonstrated, for example, by Turberfield and coworkers^30, 31^. Externally controlled, fuel-driven DNA molecular motors such as bipedal walkers suffer from two fundamental problems, however. The first is the irreversible trap-state effect, also called the “hook effect”^33^, in which two fuels of the same sequence consecutively bind the motor, one to the leg and the other to the foothold. In a properly operating walker, a single fuel strand should bind the leg and the foothold. When two separate fuels are bound to the leg and the foothold, the walker dissociates upon the introduction of the next antifuel. The second issue is the accumulation of redundant fuel and antifuel command strands and fuel/antifuel duplex waste product in the solution, leading either to walker detachment or walker striding the wrong direction. We have recently resolved these issues with microfluidics-enabled washing steps^34^ and development of an operation strategy called ‘Fuel Before Anti-Fuel’ (FBAF) that avoids the trap state, while allowing use of high fuel and antifuel concentrations that increase motor speed. With these methods we have demonstrated a bipedal walker striding on origami track for 50 steps at a 480-s stepping rate with 98% yield per step^35^.

Here, we developed a DNA origami rotary motor that, powered by DNA fuels and antifuels, operates with high processivity and speed. The rotor, which consists of a bottom stationary disk and a top rotor disk, is propelled by two DNA bipedal walkers that stride on a circular track (Fig. 1A). The bipedal walkers consume fuels and antifuels provided by a microfluidic device using the FBAF operational strategy. To prevent termination in case of operational errors (i.e., failed leg-foothold connectivity), the disks are permanently connected to each other by a single-stranded swivel element that allows free rotation of the top rotor disk while preventing its dissociation from the bottom disk^28, 29^. Thus, in contrast to linear bipedal motors that dissociate from the track in the event of an operational error^19, 35^, the connection of the disks allowed the walkers legs to rebind the track when the connection is lost and therefore fully recover and resume operation, increasing processivity.

**Fig. 1.**
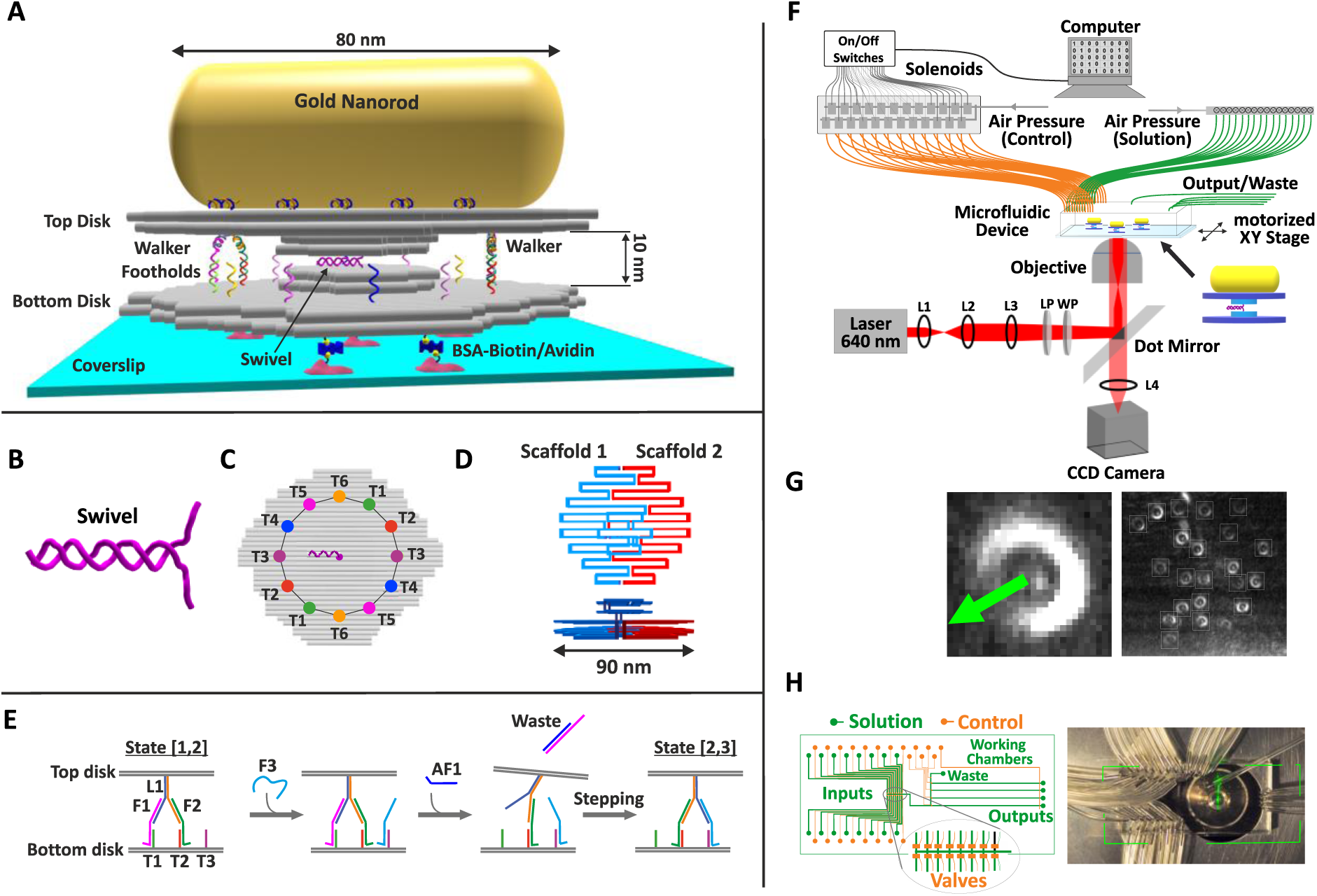
Rotor and experimental design. **(A)** DNA origami rotary motor design. **(B)** DNA swivel consisting of double-stranded and single-stranded segments. **(C)** Circular track consisting of twelve footholds denoted T1-T12. **(D)** Scaffold interlocking scheme viewed from the side (left) and top (right). **(E)** Fuel-before-antifuel walking mechanism. **(F)** Experimental setup consisting of the microfluidic system and defocusing optical setup. **(G)** Exemplary defocused image of a single gold nanorod (left) and full defocused image of ∼20 gold nanorods coupled to motors immobilized on the surface of a microfluidic device (right). Green arrows indicate rotors orientations. **(H)** Left: Microfluidic device schematic. Right: Top view of the microfluidic device placement in the optical setup.

Several light-based observation methods have been employed to monitor DNA origami rotary devices; however, various limitations rendered previously reported methods insufficient for our needs. Forster resonance energy transfer can provide information from only a limited number of states^36^, whereas our rotor assumes 24 different states and sub-states that should ideally be identified. Single-molecule total internal reflection fluorescence imaging microscopy provides high angular resolution^23, 27, 28, 29^, but this method requires a structural extension element of length around 300 nanometers to hold fluorophores away from the rotation axis (typically an origami rod with length comparable to the light diffraction limit) that typically interacts with the glass surface on which the rotor is placed, suppressing rotation^23, 27, 28, 29^. Furthermore, photobleaching and limited brightness limit resolution and measurement duration times of methods that employ organic fluorophores^36^. To monitor our rotary motor, we used single-molecule defocused orientation imaging of gold nanorods^4, 37, 38^, developed based on a previous study of the biological ATP-synthase rotary motor with high angular and temporal resolutions (∼1 degree/10 µs)^4^. The gold nanorods are highly photostable, the scattering intensity is essentially limited only by the radiation power, and the large scattering cross-section enables very good signal to noise ratio, resulting in excellent resolution^4^.

## Results

### Rotor design

The DNA origami rotary motor consists of two separately prepared but structurally identical origami disks that are connected to each other through hybridization of two swivel elements (Figs. 1A and B). Each disk is constructed from four square-lattice origami layers. The two larger diameter layers (∼90 nm) were designed to house a circular foothold track ∼60 nm in diameter (bottom disk) and two sets of bipedal walkers (top disk, ∼60 nm apart). The two smaller concentric layers (∼40 nm in diameter) were designed to maintain a gap of ∼10 nm between the disks to provide room for free operation of the walkers and to house the swivel element. The swivel was made of two elongated staples that zipped to form 30 base pairs upon interaction of the two disks, leaving 8 unpaired bases that act as the rotation axis (Fig. 1B).

To avoid collision between the disks during rotation, the disks must be as flat and as robust as possible. For robustness, the origami disks were designed with four layers, and were optimized iteratively using the mrDNA simulation package^39^ (see Supplementary Section 1 and Supplementary Fig. S1 for initial designs, iterations, and discussion of design considerations). The circular track consists of two identical sets of six footholds, ∼15 nm apart (Fig. 1C). In this design, the footholds are close enough for efficient stepping of the bipedal walkers but far enough apart to prevent unwanted crossing between sets of footholds; this design relied on our previously determined dependency of bipedal walking efficiency on step sizes^40^. This disk design requires a ∼15,500 base-pairs, twice the size of a typical origami scaffold. Therefore, each disk was assembled using two orthogonal scaffolds that interact according to the ‘scaffold interlocking’ principle in a one-pot thermal annealing process, a method recently developed by Dietz and co-workers^41^ (Fig. 1D).

To monitor our rotary motor, we imaged a gold nanorod. To achieve a strong plasmon resonance signal, the gold nanorod was ∼40 nm diameter and ∼80 nm length (640 nm wavelength^4^), slightly shorter than the origami disks to prevent nanorod interaction with the coverslip surface (Fig. 1A). Biotinylated strands, positioned under the bottom disk, were added to facilitate the immobilization of the rotor on the coverslip surface via biotin-avidin interactions (Fig. 1A).

#### Fuel-before-antifuel propulsion mechanism

The fuel-before-antifuel propulsion mechanism is based on our previous design of a linear bipedal motor^34, 35^ (Supplementary Fig. S2). This strategy avoids the irreversible trap-state effect^33^, while allowing high concentrations of fuels and antifuels for fast rotor response (Fig. 1E, see Supplementary Movie 1 for an animation of the rotary mechanism)^34, 35^. With this method, a high concentration of next-state fuels (e.g., 10 µm of F3; Fig. 1E) is introduced when both legs are still placed on current-state footholds (T1 and T2); this state is referred to as state [1,2]. The fuel quickly binds the corresponding foothold (T3), followed by microfluidics washing to remove redundant fuels (F3). These fuels cannot bind the corresponding leg (L1), which in this stage is still blocked by the previous fuel (F1), therefore avoiding the trap-state effect regardless of fuel concentrations. Antifuel strands (AF1) are then introduced at high concentrations, rapidly lifting the leg (L1) and removing the previous fuel (F1), followed by attachment of the lifted leg to the next fuel (F3) to complete the leg-placing reaction. This brings the walker to state [2,3]. This propulsion mechanism is a significant improvement over the classical operational scheme in which the leg is lifted first, followed by the introduction of fuels^35^. For limitations of the fuel-before-antifuel mechanism related to the incomplete removal of fuels by antifuels, see Supplementary Section 2.

#### Experimental setup

Our experimental setup consists of a single-molecule defocused-imaging of gold nanorods optical setup^4^, coupled with a microfluidic system^34^ (Fig. 1F, see Supplementary Sections 3 and 4 for full descriptions). Typically, 20-25 bright, defocused imaged shapes corresponding to light scattered from the gold nanorods belonging to 20-25 rotors positioned on the surface of the microfluidics glass coverslip were observed simultaneously (Fig. 1G). The imaged shapes were fitted to a set of precalculated theoretical shapes^37^ from which in-plane and out-of-plane orientations of the nanorods were determined (Supplementary Fig. S3). To facilitate stable, long-term observation, we developed real-time drift compensation and tracking procedures.

The microfluidic system allows convenient user- and computer-controlled introduction and, most importantly, removal of chemical inputs to and from the rotary motors immobilized inside the microfluidic device. The device, which is mounted on the defocused imaging setup, includes 38 input channels that convey solutions containing the molecular components necessary for rotor immobilization, assembly, and operation (Fig. 1H). The pneumatic valves that regulate flow are embedded in the device in a design that enables rapid and precise computer-controlled exchange of small-volume solutions in reproducible sequence and timing for hundreds of cycles over days of rotor operation^34^.

#### Rotor assembly

A detailed description of the rotor’s assembly is provided in Supplementary Section 5. In short, the two origami disks were separately thermally annealed, followed by purification using polyethylene glycol precipitation (Fig. 2A). The disks were then mixed in a 1:1 ratio and left to react through the swivel elements overnight (10 nM each disk, 35°C), achieving ∼70% dimerization yield, as shown by agarose gel electrophoresis (Fig. 2B, lane 3). Finally, the DNA-functionalized gold nanorods were added; the poly-T sequences on the nanorods bind with the poly-A sequences on the upper side of the top disk. Binding of the nanorods was confirmed by agarose gel (Fig. 2C, lane 2). That the desired products of the stepwise process were obtained was confirmed by transmission electron microscopy (TEM) (Fig. 2A, D). Additionally, toehold-mediated swivel displacement demonstrated the expected rotor disassembly (Fig. 2B, lane 4, and 2C, lane 3), confirming that the interaction was solely through the swivel.

**Fig. 2.**
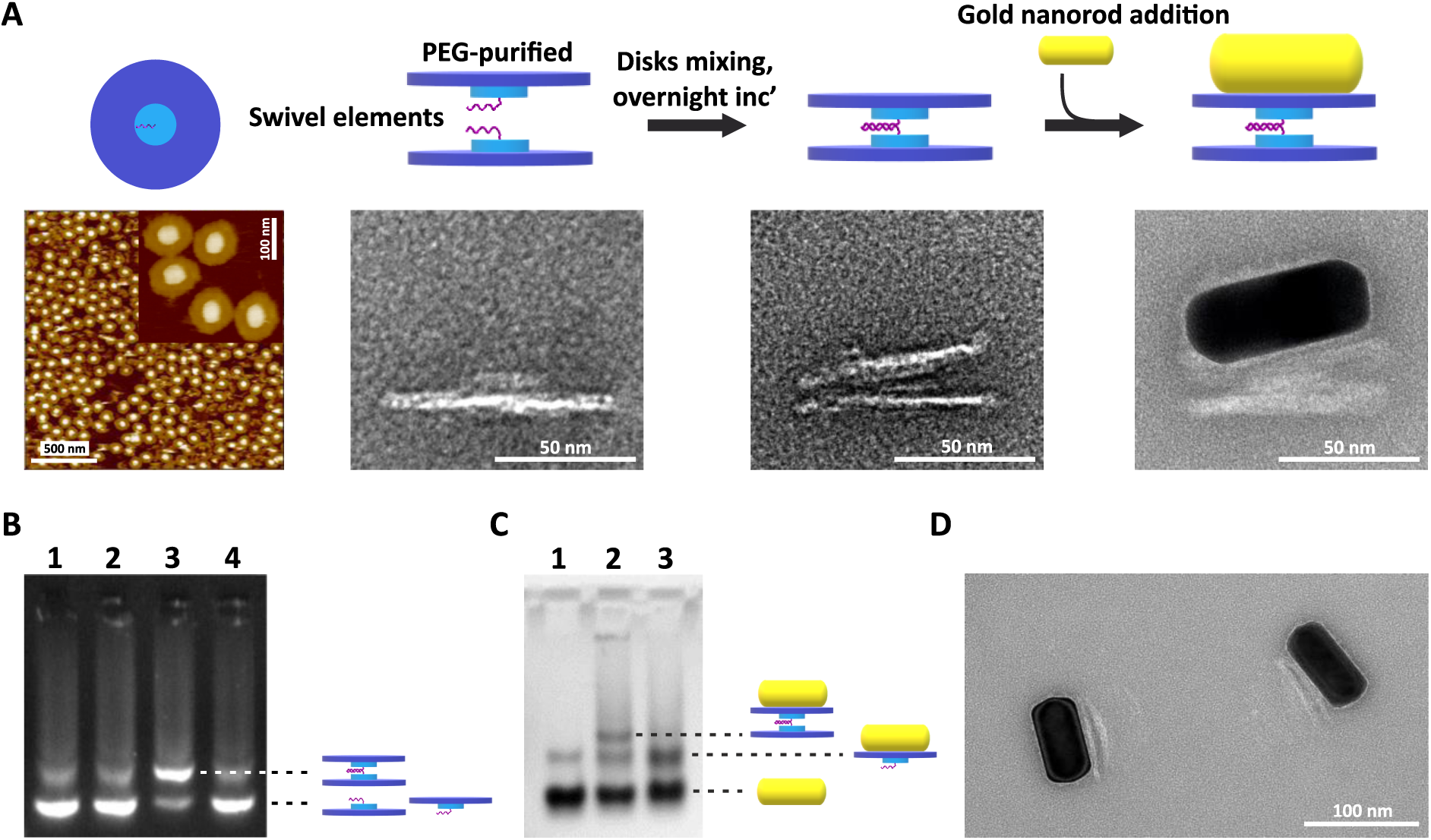
Rotor assembly. **(A)** Schematic of assembly process and images of intermediates and products from left to right: AFM images of the disk monomer, TEM image of the disk monomer, TEM image of the rotor (i.e., disk dimer), and TEM image of the rotor coupled to a gold nanorod. **(B)** Agarose gel analysis of the disk dimerization reaction. Lane 1: top disk monomer; lane 2: bottom disk monomer; lane 3: rotor; lane 4: dissociated rotor. Dotted lines indicate disk monomer and dimer bands. **(C)** Agarose gel analysis of the nanorod coupling reaction. Lane 1: gold nanorod; lane 2: rotor mixed with gold nanorod; lane 3: dissociated rotor and gold nanorod. Dotted lines indicate gold nanorod, top disk coupled to a gold nanorod, and rotor coupled to gold nanorod bands. A small population of gold nanorod dimers travels at the same speed as top disks coupled to gold nanorods. **(D)** TEM image of rotors coupled to gold nanorods extracted from the agarose gel (panel C, lane 3).

#### Rotor operation

To immobilize the rotors inside the microfluidics device, a solution containing the fully assembled rotors (∼0.2 nM) was introduced into the working chamber of the device until the desired density of rotors was immobilized on the coverslip surface through a biotin-avidin interaction (Fig. 1G). This typically required several minutes. Initially, rotors are in Brownian rotation mode as fuels are not introduced during this preparation (for a characterization of the Brownian rotation state, see Supplementary Section 6 and Supplementary Fig. S4). To initialize the rotors for directional rotation, the bipedal walkers were attached to the track at state [1,2] using fuels and antifuels (the track binding reactions and preparation procedure for directional rotation are described in Supplementary Figs. S5 and S6, respectively).

Once initialized, the rotors were commanded to perform a full 360° rotation consisting of 12 steps between 12 distinguished states using the FBAF procedure. This required 24 consecutive leg-lifting and leg-placing reactions (Fig. 3A and Supplementary Movie 2). Examples of rotors performing 4, 5, and 8 full rotations, corresponding to 48, 60, and 96 rotor steps, respectively, are shown in Fig. 3B and Supplementary Movies 3-5. Rotors often operated correctly over days; however, close examination of the rotation time-traces revealed short episodes of leg dissociations that resulted in free Brownian rotation or slow rotor response to commands, followed by rotor recovery and return to normal operation (Fig. 3B, see also Fig. 4D). Rotors operated at different speeds (i.e., different command strand introduction rates) and rotors walking in alternating directions (i.e., different command sequences) are shown in Fig 3C and 3D, respectively, and in Supplementary Movie 6.

**Fig. 3.**
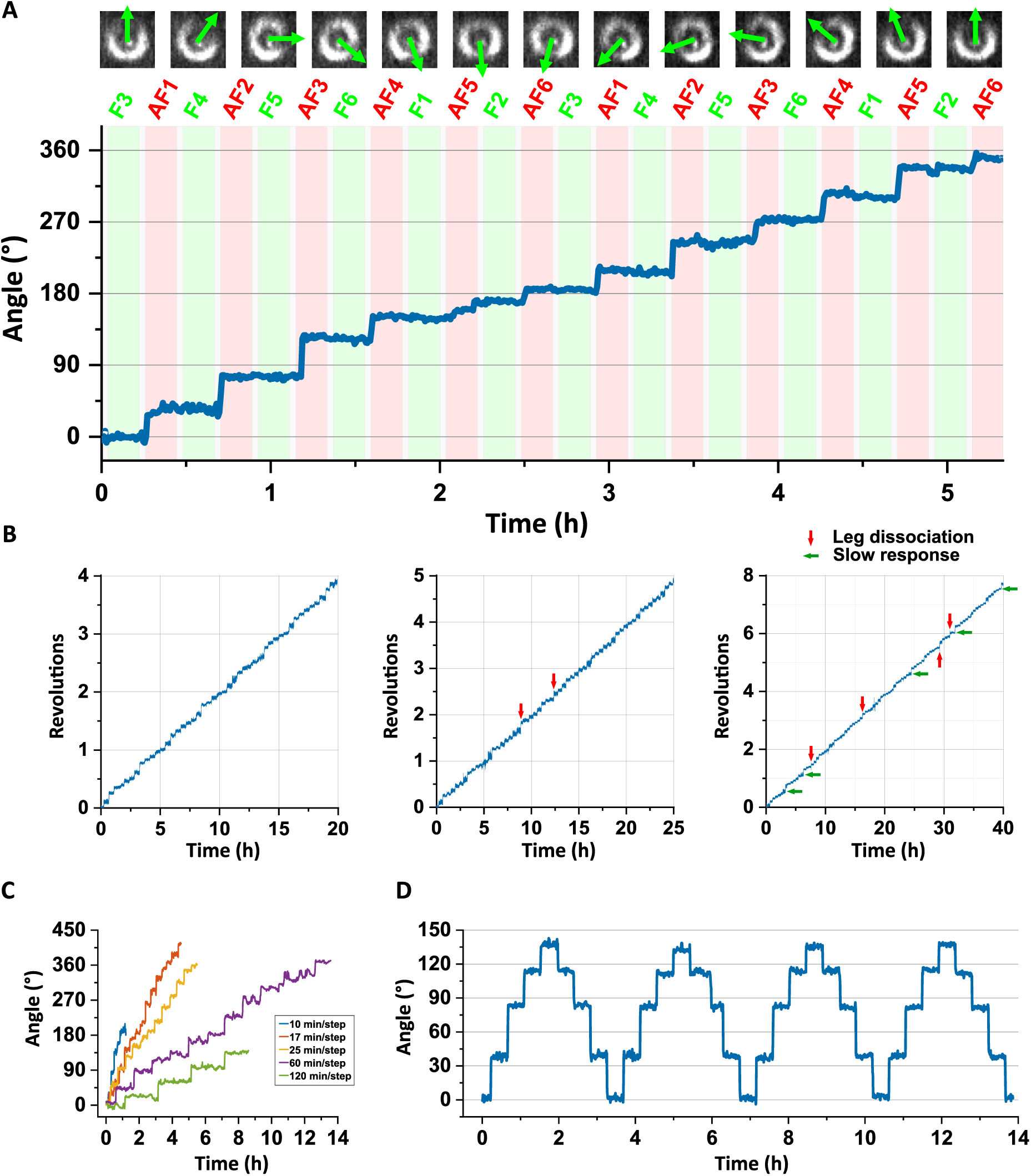
Rotor operation. **(A)** Rotor performing 12 clockwise steps consisting of a full revolution. State [1,2] is the initial and final state. Top: Defocus shapes between each of the steps. Bottom: Angular time-trace of the rotation. Incubation periods of the fuel and antifuel commands are indicated by green and red backgrounds, respectively. Washing periods are indicated by a grey background. Bin size of 50 data points. **(B)** Clockwise revolutions as a function of time for three exemplary rotors. Leg dissociation and recovery events and slow rotor responses are indicated by red and green arrows, respectively. **(C)** Rotor angle as a function of time at five different computer-controlled command strand introduction rates. Bin size of 100 data points. **(D)** Rotor angle for rotor rotating in alternating directions. Bin size of 100 data points.

**Fig. 4.**
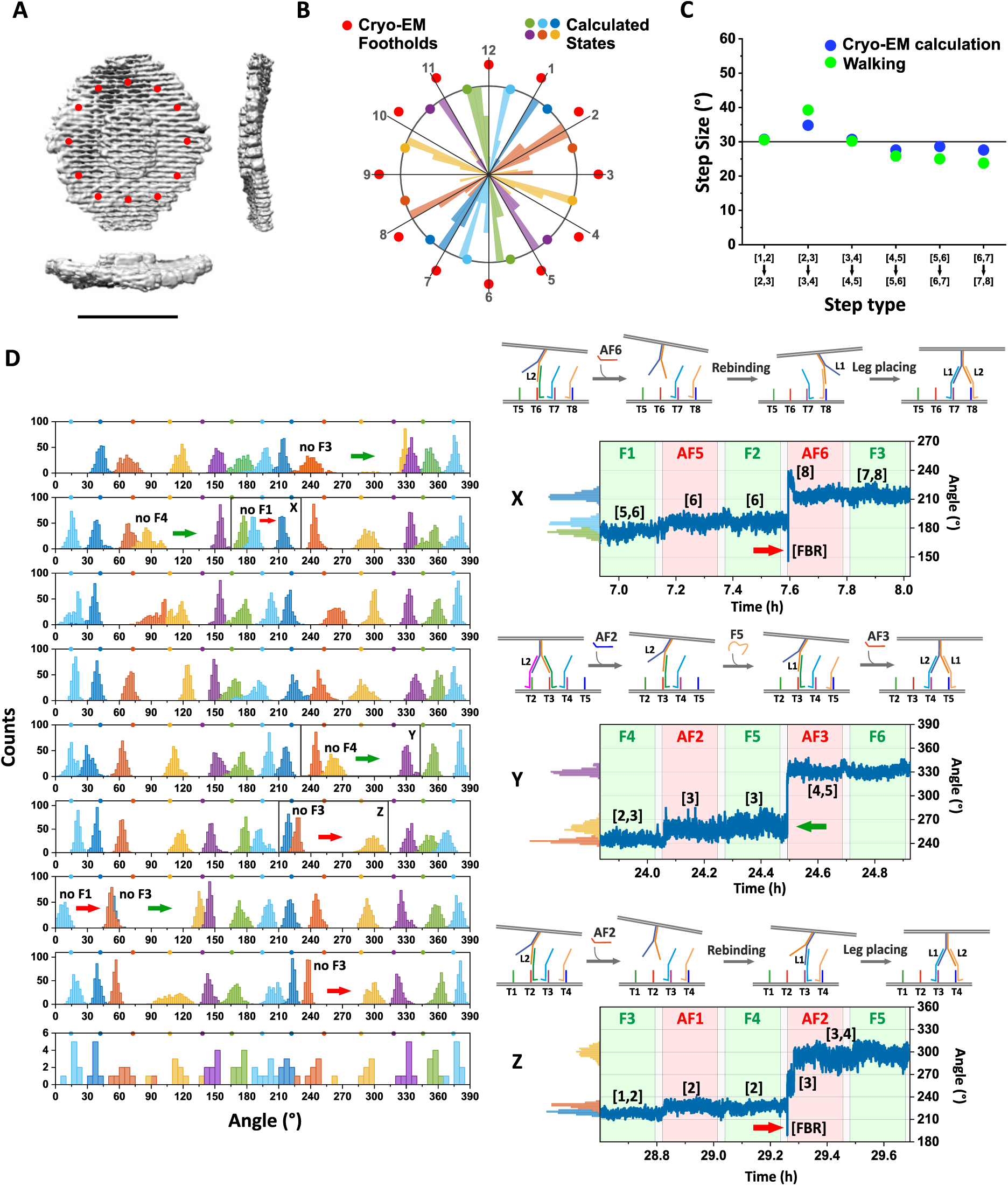
Structure of the track. **(A)** Cryo-EM 3D model of the origami disk shown in top and side views. Foothold positions are indicated by red dots. Scale bar: 50 nm. **(B)** Comparison of cryo-EM-based foothold positions (red dots), calculated rotor states (middle positions between footholds), and histograms of measured rotor states (obtained from the right-most rotor in Fig. 3B). Each of the 12 histograms consist of 8 data points averaged from the angular trace during fuel-incubation periods. **(C)** Comparison of cryo-EM-based step sizes and measured step sizes (∼120 steps for each of the six step types). **(D)** Left: Rotor orientation histograms measured at 96 different states throughout 8 rotor revolutions (same rotor as in panel B). The data used were extracted from fuel incubation periods and were binned with a bin size of 10. Red and green arrows indicate errors of leg dissociation and slow rotor response, respectively. Missing fuels prior to errors are written next to the corresponding states. The bottom panel summarizes the average positions for each of the 12 states. Right: Three exemplary errors denoted cases X, Y, and Z. The interpretations of rotor operations are illustrated above the time traces, and the corresponding rotor states are displayed along the trace. FBR denotes free Brownian rotation. Red and green arrows indicate errors of leg dissociation and slow rotor response, respectively.

#### Structure of the circular track

Figure 4A shows the Cryo-electron microscopy (cryo-EM) reconstructed model of the origami disk. The red dots indicate the designed positions of the 12 footholds, designed to be located at ‘round-hour’ orientations as in a dial clock (Fig. 4A). Limitations in staple positions and minor overstretching of the origami structure along the vertical axis resulted in some of the footholds diverting by up to several degrees from the ‘round-hour’ orientations (Fig. 4B). The 96 experimental rotor state orientations were measured for an example rotor (Fig. 4B, same rotor as in Fig. 3B, right most time trace). There was general agreement between the expected and the measured orientations, with a maximum error of 14° in the case of state [2,3]. In addition, there was general agreement between the cryo-EM-based expected step sizes and the measured averaged step sizes (Fig. 4C). For example, the steps sizes around 3 and 9 o’clock were slightly larger than 30 degrees, and the steps sizes around 6 and 12 o’clock were slightly smaller than 30 degrees.

#### Rotor operational errors and rotor recovery

Rotor orientation histograms were measured at 96 different states (8 full rotations), with most of the histograms indicating the correct orientation, and a small minority of histograms are as far as 45 degrees from the expected value (Fig. 4D, left). Examination of the orientation time traces revealed episodes of dissociation of all four legs and free Brownian rotations, and of slow rotor response to external antifuel commands (Fig. 4D, right).

For example, in case X, an episode of rotor free rotation was clearly detected after introduction of antifuel AF6 (Figure 4D, right), which indicates that leg L1 did not properly connect to foothold T7 (via fuel F1) in the preceding step. This conclusion is further supported by the fact that the rotor barely responded to the introduction of antifuel AF5 in the preceding step, indicating lack of proper leg L1 attachment to foothold T1. Upon the introduction of antifuel AF6, the rotor dissociated from the track for about 300 milliseconds, followed by placement of the leading leg L2 on T8, followed by placing of the trailing leg L1 on T7 about 60 seconds later, completing recovery to normal operation. A similar mechanism of recovery was observed in case Z, but with the trailing leg L1 binding first to foothold T3, followed by binding of the leading leg L2 to T4 (Fig. 4D, right). More cases of rotor dissociation followed by fast recovery are shown in Supplementary Fig. S7.

In another type of recovery, case Y, the rotor barely rotated forward upon the introduction of antifuel AF2, as is in cases X and Z, but no dissociation was observed upon the introduction of subsequent antifuel AF3 (Fig. 4D, right). In such cases, we cannot determine whether the rotor dissociated as in cases X and Z, but rebinds quickly within the measurement time resolution (300 milliseconds) or whether the leg placing mechanism worked properly, and only the body of the origami rotor was slow to respond.

#### Rotor speed and response rate

In externally controlled motors, such as the rotor presented here, motor speed is determined by the rate of command strand introduction (Fig. 3C). High rates of introduction of fuel and antifuel strands result in rapid rotation but shorten the time allocated for the leg placing and lifting reactions, which may result in rotor malfunction. Thus, high intrinsic leg placing and lifting reactions rates are necessary for fast and processive rotors.

Kinetic analysis of the stepping reactions initiated by the introduction of the antifuels based on averaged rotation measured for ∼120 steps (for each of the six different antifuels, taken from 12 different rotors) do not show a single exponential response and a completion of the reaction within a given period, but instead show two phases, including a fast response for most of the rotors and a slow, or no response for the others (Fig. 5A). Attempts to fit these data with a double exponential function, describing fast and slow responding populations, or a double exponential function representing a reaction with a stable intermediate, describing leg lifting followed by leg placing, were only partially successful (for additional discussion, see Supplementary Fig. S8).

**Fig. 5.**
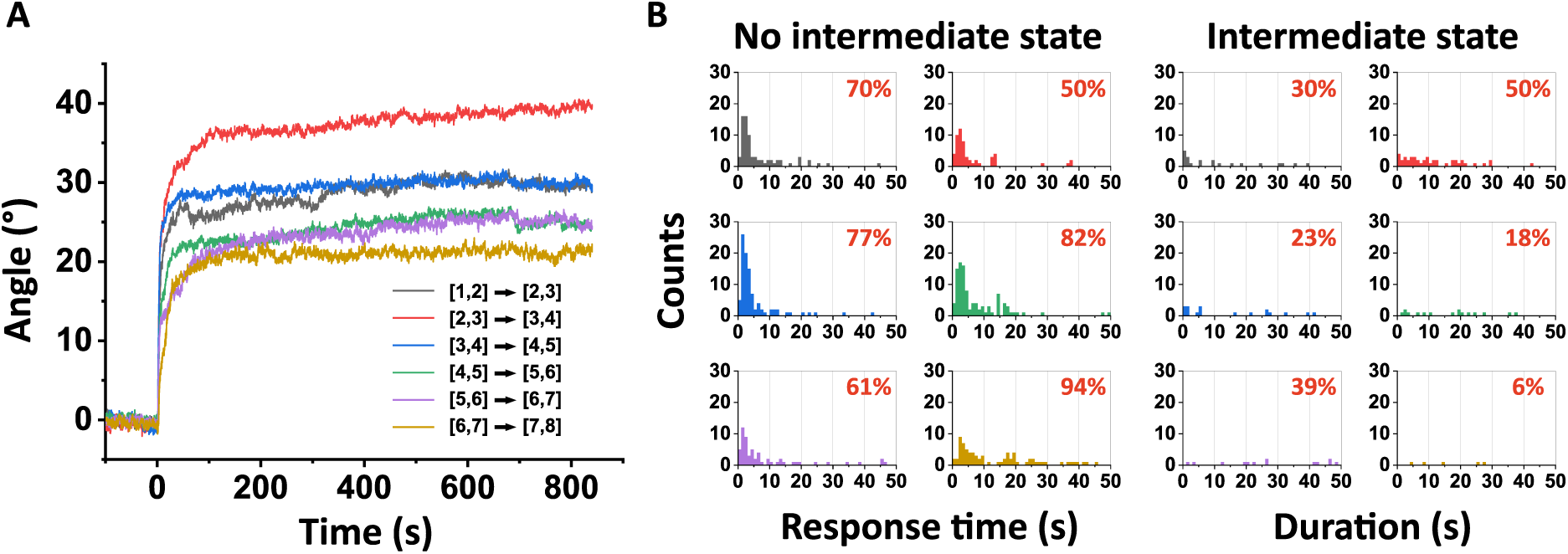
Motor speed. **(A)** Averaged angles as a function of time for ∼120 steps for each of the six step types. **(B)** Histograms of rotor response times extracted from the step time-traces (same dataset as in panel A). Left: response time in cases where no intermediate state was detected. Right: Duration of detected intermediate states. In both cases only the first 50 seconds are shown, but small “tails” of response times and intermediate state durations of up to 600 seconds were also observed and were considered in the displayed percentages.

Therefore, instead of fitting these ensemble-level kinetic profiles, individual steps were fitted using a single-step function, representing an immediate transition (within the resolution of the measurement) or a two-step function, representing such a transition but with an apparent intermediate state (see Supplementary Fig. S9 for exemplary fitted steps). The former are cases in which the rotor completes the transition in one apparent step (Fig. 5B, left), and the latter are cases in which the rotor shows an intermediate state (e.g., the leg is lifted but not yet placed, Fig. 5B, right panels).

For most of the stepping reactions, an intermediate state was not observed (Fig 5B, percentages in red), and for most of these cases, the stepping reaction was completed within 1 to 7 seconds after introduction of the antifuel. A minority of the rotors responded more slowly (from 10 to more than 100 seconds), indicating difficulties in completion of the stepping reaction, as also indicated by the presence of the slow components in the kinetic profiles (Fig. 5A, see also Supplementary Fig. S10A for a zoom-in on the normalized profiles). The time it takes for the microfluidics to deliver the antifuel commands plus the antifuel binding time is about ∼3 seconds, resulting in a corresponding delay in rotor response (Fig. 5B, left). Furthermore, in cases in which an intermediate state was observed (Fig. 5B, right), the intermediate state duration was 10 to more than 100 seconds. In addition, the kinetic profiles revealed significant differences in the response times of the six different step reactions. For example, the step [3,4] → [4,5] is, on average, ∼10 times faster than the step [5,6] → [6,7] (for details see Supplementary Fig. S10B-D). Taken together, our kinetic data shows that for most steps and most rotors the stepping reactions are completed within less than 10 seconds, representing properly functioning rotors. However, occasionally, stepping would not take place before the subsequent antifuel is introduced, resulting in walkers dissociation.

#### Rotor limitations and possible improvements

Our analysis showed that for most steps and most rotors the stepping reactions were completed within less than 10 seconds, but for a considerable minority of cases, stepping reactions required hundreds of seconds (in a non-single exponent or a non-first-order fashion). Further, the data showed significant differences in rotor response in terms of yields and speeds between the six different commands. In principle, two types of rotor improper operation explain these results. One possibility is malfunctioning of the propulsion mechanism, that is, improper interaction of the legs, fuels, antifuels, and footholds. Another possibility is rotational hindrance caused by unwanted interaction of the upper origami or the gold nanorod with the bottom origami or the surrounding coverslip.

To minimize the amount of rotational hindrance experienced by our rotors, we used a working buffer with minimal concentrations of cations (Supplementary Fig. S4A, B). Under these conditions, although some freely rotating rotors (in the absence of fuels) were stuck for up to 3 minutes, rarely were rotors stuck for 10 minutes, the time window we provided the rotors to respond during operation (Supplementary Fig. S4C, D). Therefore, most of the cases of slow responding or non-responding rotors that led to dissociation were not due to stuck rotors.

Nevertheless, redesigning the rotor such that the gold nanorod is smaller than the upper disk, which in turn is smaller than the bottom disk, should prevent unwanted interactions between the nanorod and upper disk with the coverslip environment, further increasing free rotation (Supplementary Fig. S11A).

Our analyses also suggest an explanation for the occasional malfunctioning of the propulsion mechanism. That there are differences in kinetic data between the six different steps indicate that the differences are not due to the positions of the footholds on the origami but rather to the sequences of the footholds, legs, fuels and antifuels involved (Supplementary Fig. S10C, D). This suggests that at least some of our sequences may be sub-optimal and can be improved significantly. Unoptimized sequences may result in the formation of secondary structures that slow, or even prevent, desired interactions. The sequences that were used for our rotors are essentially identical to those used in the original bipedal motors developed by Shin and Pierce^42^, with some optimization for minimum secondary structures. Our data suggests that more comprehensive optimization may result in better operation.

Further, the fuel-before-antifuel operational mechanism, despite its benefits, has one major drawback: the formation of an unwanted heterocomplex between the trailing leg antifuel and the leading fuel that prevents the completion of the stepping reaction^35^ (for details see Supplementary Fig. S2A). These complexes are typically removed by thermal dissociation, but in some cases, they remain bound, slowing rotor response. Such interactions are unavoidable, but negative influences may be mitigated by the use of two short antifuel strands instead of one long antifuel as used here, that may lead to exponentially faster heterocomplex removal (Supplementary Fig. S2B).

Another possible detrimental effect is molecular threading of the legs, footholds, or the foothold-fuel complexes into the origami disks (Supplementary Fig. S11B). This interaction would render the strand unavailable to execute the propulsion reaction (for specific examples, see Supplementary Fig. S12). Such unwanted interactions may be reduced by distancing the footholds and the legs from the origami body by a supporting strand elongated from the neighboring staple. Alternatively, an additional identical parallel foothold-track and additional pairs of legs could compensate for unavailable footholds and legs and should handle cases of truncated fuel strands better than a single system (Supplementary Fig. S11C).

### Conclusions

In conclusion, we presented here an externally controlled DNA origami rotary motor in which a top disk is propelled around a bottom disk by two bipedal walkers that consume DNA fuels. To facilitate high processivity, we employed several strategies: computer-controlled microfluidic operation, an operational mechanism free of trap states, and a swivel connection between the disks that allows the rotor to recover from operational errors and returning to normal operation. Our results show that these strategies were successful: The rotor demonstrated 8 clockwise revolutions – nearly 100 steps, rotation in alternating directions, a common feature in macroscopic rotary motors, and we demonstrated excellent computer-control over nanomotors.

Here we used single-molecule defocused imaging of gold nanorods to optically monitor rotary motor function. This strategy allowed us to observe the rotor’s Brownian rotation, track binding reactions and stepping rotation. The high angular resolution of the method enabled us to clearly resolve small rotational transitions of several degrees. The high time-resolution of the system resolved step completion times of around a hundred steps in a single experiment with sub-second resolution. Importantly, very long measurements were enabled by the stability of the nanorods; rotors were monitored for several days with no degradation in signal quality. Additionally, these measurements allowed us to follow hundreds of chemical reactions administered by the microfluidic device. We expect that the methods employed here, whether combined or separate, will be valuable for investigations of molecular rotary systems.

A natural next step will be to develop an autonomous rotary motor that runs on DNA fuels. Autonomous mechanisms based on DNA fuels have been described, although not implemented for more than two steps due to a lack of a robust track^30, 31^. The externally-controlled DNA origami rotary motor developed here could therefore serve as a reliable structural platform for the implementation and development of much sought after autonomous mechanisms.

## Acknowledgements

This work was supported by the Israel Science Foundation. We thank Narain Karedla and Joerg Enderlein for assistance in defocused imaging optics and analysis software, Hiroyuki Noji for assistance in defocused imaging optics and dot mirror preparation methods, Enzo Kopperger and Friedrich Simmel for providing DNA origami rotor prototypes, Thomas Gerling and Hendrik Dietz for assistance in 3D origami design, Doron Gerber and Noa Marchoom-Bura for providing microfluidic devices, Massimo Kube for assistance in Cryo-EM characterization of origami structures, and Ran Zalk for cryo-EM measurements and analysis.

## Data and Code availability

Data in this paper as well as the cadnano file of the DNA origami Disk are available upon request. The custom software used for defocused imaging data analysis can be found in <u>github</u>.

## Supporting Information

## Section 1: DNA Origami Design

### Design of the DNA origami disk

The DNA origami disk was designed as a square lattice^1^ using cadnano 2.0 (Fig. S1A-C). The scaffolds used were CS4 of length 7758 and P8064 of length 8064, both purchased from Tilibit Nanosystems. Simulations were performed using MRDNA software^2^.

### Disk twist minimization

Although the global twist that results from the non-integer number of bases between crossovers can theoretically be corrected by periodic deletion every 64 bases, in some cases there is a residual global twist^2^. To minimize this twist, an iterative simulation process of various origami designs and deletion methods was performed^2^. Our initial failed designs that contained significant twists consisted of a single-layered disk, single-layered disk with periodic deletions every 32 or 64 bases, and a double-layered disk with perpendicular layers^3, 4^ (Fig. S1D). Our final design, which was sufficiently flat, was double-layered and contained deletions every 32 bases.

### Scaffold interlocking

A scaffold interlocking strategy^5^ facilitated construction of the large and multi-layered disk structure. To minimize incorrect interactions, we selected the highly-orthogonal scaffolds P8064 and CS4 (Tilibit Nanosystems). In our design, the interlocking segments were 100 bases long and had similar GC contents. Due to its lower reliability, an interlocking region was not incorporated in the sensitive inner part of the disk where the two disks are near each other and might interact if not properly sealed (the inner part consisted of only P8064, Fig. S1E). Additionally, special attention was paid to sequences that were homologous between scaffolds (with maximum lengths of 89 and 107 bases) and within the P8064 scaffold (with a maximum length of 42 bases). To minimize the effect of possible incorrect interactions involving these sequences, they were positioned such as to avoid the scaffold interlocking regions and the specific staples that are extended with rotor machinery (e.g., legs, footholds).

### Staple design

For the DNA origami disks, we used staples with lengths of 48 to 64 bases, a recommended length^2^. We maximized crossover density and maintained a presence of at least one continuous segment of 16 bases for each staple^6, 7^. Longer staple lengths are less reliable due to a lower yield of full-length DNA strands when produced in the common solid-phase synthesis techniques such as those used by IDT. To maximize the available positions for rotor machinery in the inner part of the rotor so as to increase design flexibility, the continuous 16 base segments were mostly placed in the outer part of the disk.

### Edge design

The disks were designed to avoid inter-origami interactions that arise from blunt-end stacking between helix edges. In addition to the typical poly-C overhangs and loops that are placed at the edges of the origami helices wherever possible (poly-C was selected over poly-T to avoid interfering with the poly-A\poly-T origami nanorod interactions), the circular design of the origami and an asymmetry between both of its sides minimized possibilities of helix alignment.

**Figure S1.**
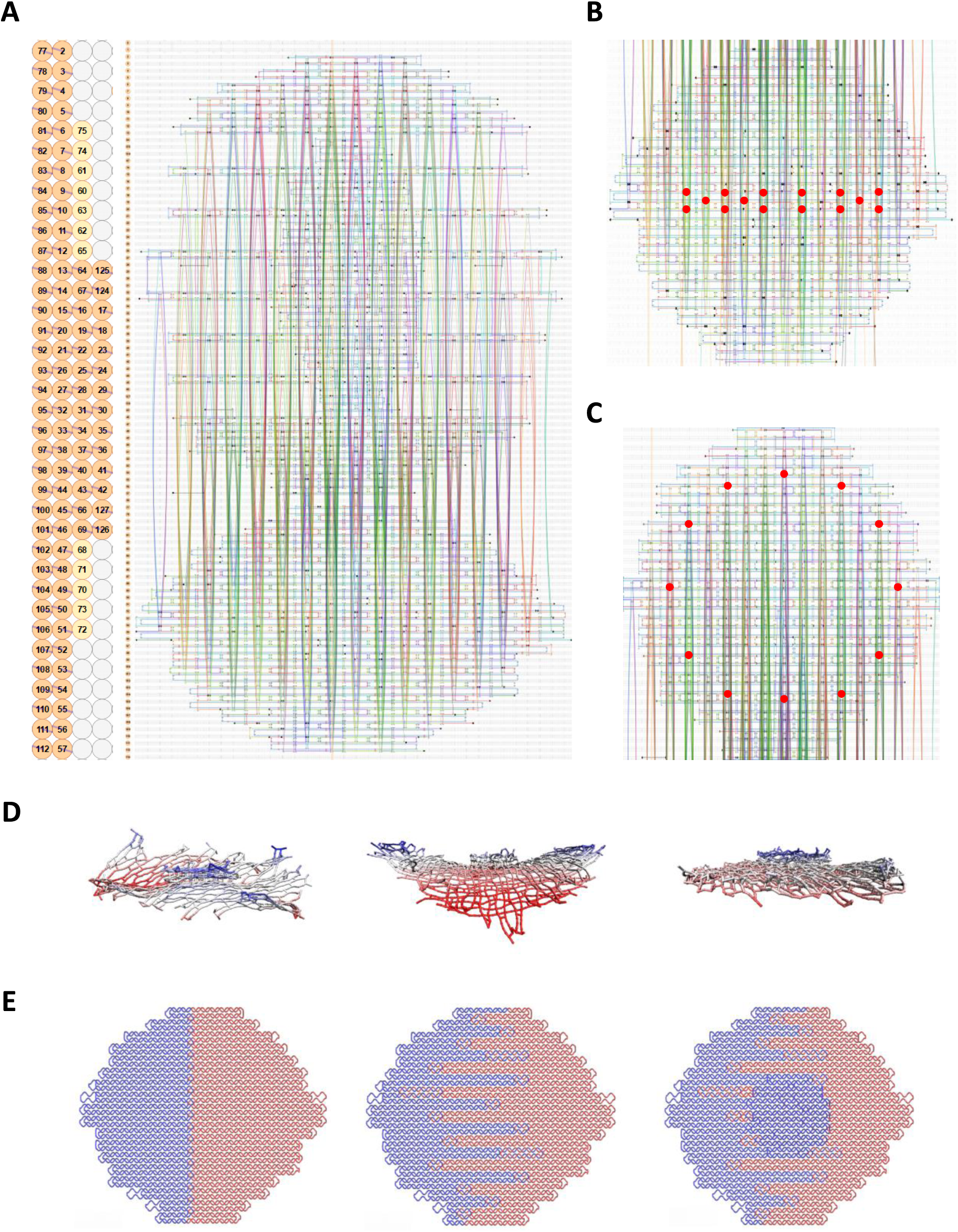
DNA origami disk design. **(A)** Full Cadnano design. **(B)** Gold nanorod binding positions on the outer side of the top disk. **(C)** Foothold placement on the inner side of the bottom disk. **(D)** Design iterations. From left to right: single main layer, two perpendicular main layers, and two regular main layers. **(E)** Scaffold interlocking scheme. P8064 and CS4 scaffolds are colored blue and red, respectively. The pictures show the bottom main layer with no interlocking segments (left), top main layer that includes interlocking segments (middle), and the full design including the central layers that belong exclusively to the P8064 scaffold.

## Section 2: Rotor Operation Mechanism

In the fuel-before-anti-fuel (FBAF) mechanism^8^ (Fig. S2A), the antifuel of the trailing leg (e.g., AF1) is inserted while the next-state fuel (F3) is attached to the foothold (T3) but not to the corresponding leg (L1). Because F1 and F3 share a common sequence that binds them to L1, the complementary sequence is shared in AF1 and AF3. This leads to formation of a heterocomplex structure between AF1 and F3 upon insertion of AF1, and a blockage of the leg placing reaction (HC1 in Fig. S2A). To avoid this blockage, in the FBAF strategy the antifuels are shortened in the segment that is involved in the heterocomplex (to 8 base pairs) such that the interaction is reversible at room temperature, allowing the heterocomplex to dissociate and the leg placing reaction to complete.

The dissociation rate of the heterocomplex was found to be a limiting factor for the stepping reaction in our study of the FBAF mechanism^8^ and could also explain the slow steps performed by the rotary motor presented here. For a faster heterocomplex thermal dissociation rate, we suggest use of two even shorter antifuels (based on 5 base pairs in the thermally dissociating segments), allowing exponentially faster dissociations (Fig. S2B).

**Figure S2.**
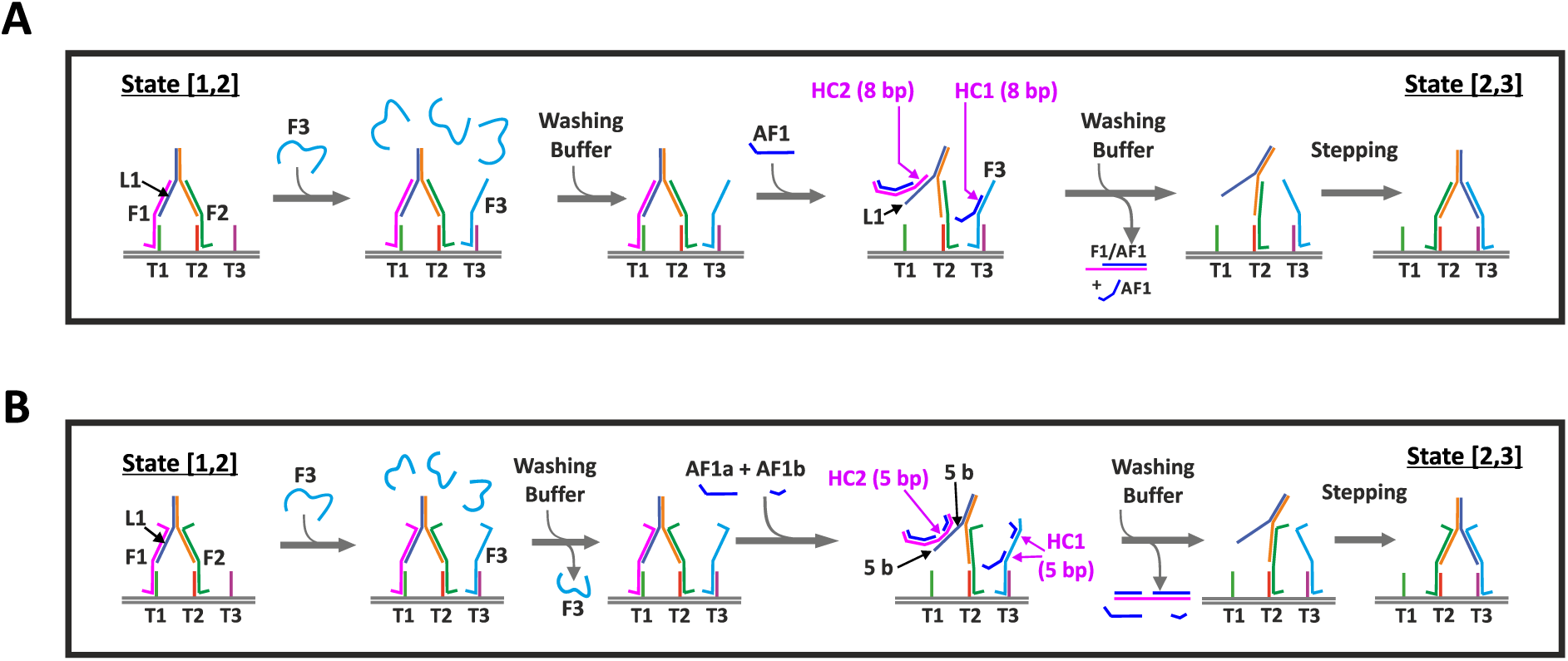
Fuel-before-antifuel operation mechanism and potential for optimization. **(A)** FBAF mechanism shown for the linear bipedal walker^8^: AF1 (blue) binds to F3 and forms a heterocomplex (HC1). An additional heterocomplex (HC2) forms between AF1 and F1, that is also 8 base-pairs in length and thermally dissociates. **(B)** In the fuel-before-two antifuels mechanism, two antifuel strands (AF1a and AF1b) separately remove the leg-fuel connection and the foothold-fuel connection.

## Section 3: Defocused Imaging

### Optical setup

Defocused imaging was carried out using a custom-built wide- and dark-field optical setup adjusted from Enoki et al.^13^ (Fig. 1F). A red diode laser (640 nm, Cube 640-40C, Coherent Europe) was passed through a single-mode fiber, collimated, expanded (measured power was ∼130 µW), reduced in diameter to remove background, focused (25-cm lens), circularly polarized, reflected by a dot mirror (fabricated in-house) toward the back focal plane of an objective (Plan Apochromat, 60x/1.45, Olympus America) mounted on an inverted microscope (IX71, Olympus America). The scattered light was collected by the same objective, passed through the glass the dot mirror around the dot (background is removed by the dot), magnified by 1.6x, and imaged onto an EMCCD camera (IXON DU-897E, Andor). For defocusing, the position of the objective was lowered by ∼1.1 µm from the focus position using a piezo scanner (P-721 focus scanner, Physik-Instrumente). The microfluidic device coupled to a glass coverslip was placed on the objective and held by a motorized XY stage (XY-MR-50-150, Holmarc).

### Dot mirror fabrication

Dot mirrors were produced in-house by methods of lithography. The dot mirror consisted of a transparent rectangle-shaped glass (25 x 36 mm) with a silver elliptic mirror (effective diameter of 3.5 mm) in its center. The layers of the ellipse consisted of a TiO_2_ adhesion layer, a silver layer (150 nm), and a silica anti-oxidation layer (50 nm).

### Data acquisition

Movies were acquired in a custom-built live-analysis pipeline (Matlab). Frames consisting of 512 x 512 pixels were acquired with an exposure time of 1 ms, and a cycle time of 80 ms for short movies or 300 milliseconds for long movies. The short exposure prevented smearing of defocused shapes that could happen during the Brownian rotation of the rotor, and the long cycle time supported long recordings, up to days. Molecule finding was performed in the focus position. The acquisition software (Matlab) was synchronized with the microfluidic command timeline through communication with the custom LabVIEW program that controlled the microfluidics.

### Fitting defocused shapes

Each observed rotor was assigned a region of interest of 29 x 29 pixels that was fitted to a set of theoretical defocus images using a least squares algorithm (Fig. S3). The theoretical images were customized to experimental conditionssuch as magnification and pixel size (using code supplied by the Enderlein group, private communication), with dot mirror optics also considered. The free parameters during the fitting were x, y, z, θ, and φ, where x and y denote the position of the region of interest, z is the defocus distance (monitored with a resolution of 0.1 µm), and θ and φ specify the orientation of the nanorod in 3D (monitored with resolutions of 10° and 1°, respectively). All orientation time-traces in this report refer to φ orientation. To resolve the inherent ambiguity of the signal to a rotation of 180°, after the first frame, the orientation was selected based on higher proximity to the previous frame. θ was constrained between 70°-90° for better fitting. To facilitate fast online analysis of up to ∼35 particles with 80 ms per frame, the fitting was done using a custom search algorithm. For each fitting, first the z and φ parameters were found roughly, and a local search was used to optimize the x, y, z, θ, and φ parameters. Following an initial calibration step performed in the first seconds of the recording, the x and y parameters were optimized only every 60 seconds.

### Observational stability

To maintain observational stability and account for drift, the x and y positions were tracked by the software, and the z position was stabilized by the objective holder in a feedback loop with the average fitted defocus distance.

#### Scanning system

For datasets containing multiple areas, areas were first collected and then scanned again, possibly between chemical reactions. The x, y, and z positions of each area and rotor within the area were maintained by the software and were used in combination with image analysis procedures to ensure correct molecule observations.

**Figure S3.**
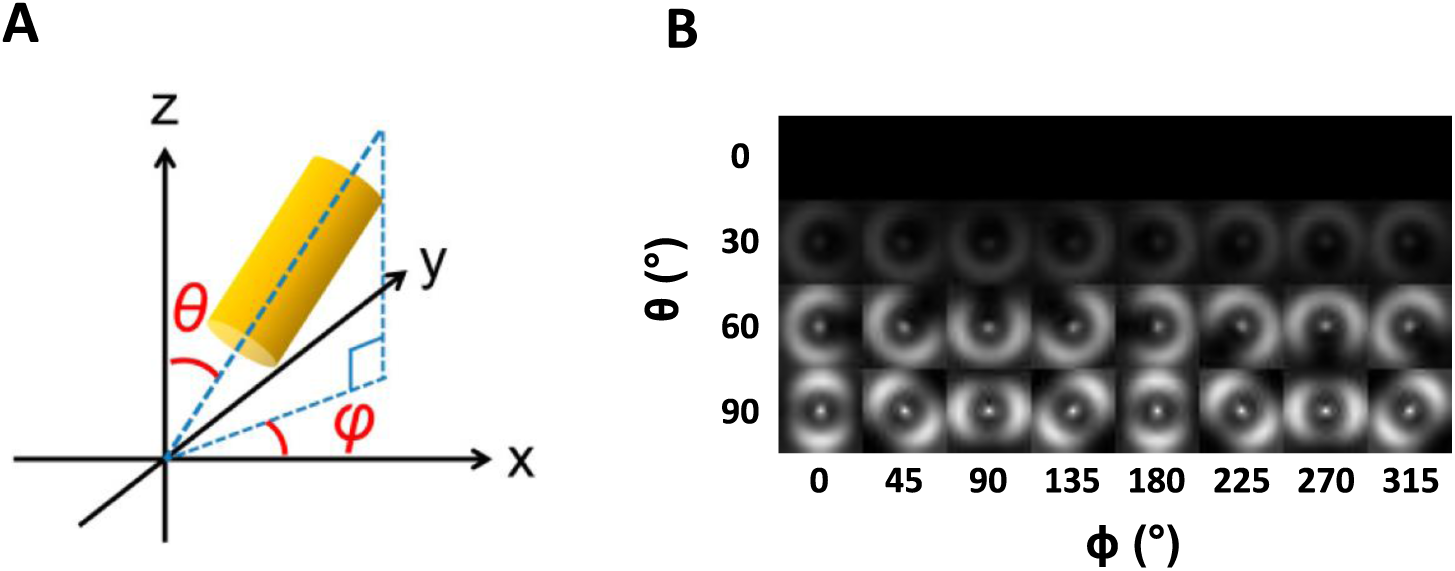
Defocused imaging. **(A)** gold nanorod orientation (illustration reproduced from Sawako et al.^8^). θ denotes in out-of-plane orientation, and φ denotes the in-plane orientation. **(B)** Theoretical defocused shapes as a function of θ and φ. The signal vanishes as θ approaches 0 due to reduced nanorod excitation. Rotor observation was not affected by this problem as rotors were typically placed straight on the surface.

## Section 4: Microfluidic Device

The operation principles and fabrication procedure of the silicone elastomer polydimethylsiloxane (PDMS) microfluidic device were previously described^14^. Briefly, the device is made of two layers: a “flow” layer that contains the channels that direct the solutions, and a control layer that contains the channels that provide pressure to “Quake-type” pneumatic valves^15, 16^. The valves allow selective flow inside the device and are computer-controlled through external solenoids using a custom interface (LabView). Device preparation included separate preparation of the polydimethylsiloxane flow and control layers, aligning and attaching them to each other through baking (80 °C), and attaching the device to a pre-cleaned glass coverslip. For cleaning, the coverslip was washed with Alconox, KOH, and methanol steps, with water rinsing in between, and both the device and the glass coverslip were subjected to an oxygen plasma treatment.

## Section 5: Rotor Assembly

### Buffers

- FOB15 (Disk folding buffer): 5 mM Tris, 1 mM EDTA, 5 mM NaCl, 15 mM MgCl_2_.
- PEG precipitation buffer: 1x TAE, 500 mM NaCl, 15% PEG 8000.
- 106 (Disk and rotor storage buffer): 1x TAE, 100 mM NaCl, 6 mM MgCl_2_.
- 102 (Rotor operation buffer, gold nanorod storage buffer, and gold nanorod coupling reaction buffer): 1x TAE, 100 mM NaCl, 2 mM MgCl_2_.
- T50 (Glass surface pre-coating buffer): 10 mM Tris, 50 mM NaCl.
- 502 (Rotor surface binding buffer): 1x TAE, 500 mM NaCl, 2 mM MgCl_2_.

### Folding the bottom and top DNA origami disks

The top and bottom disks were folded separately. Each mixture contained FOB15, 20 nM of each of the P8064 and CS4 scaffolds (Tilibit Nanosystems), and 200 nM staples (IDT). The concentration of the swivels, biotin anchor hybridization staples, and biotinylated anchor strands was 60 nM. The folding solutions were annealed in a Biorad thermocycler starting with heating in 65 °C for 15 min and subsequently reducing the temperature from 60 °C to 44 °C at a rate of 1 °C/h. The folded structures were stored at room temperature until further use.

### Connecting the disks (disk dimerization)

The folded top and bottom disks were purified using a PEG precipitation protocol described by Wagenbauer et al.^9^ with minor modifications. There were two rounds of precipitation, where the origami solutions (50 or 100 µl) were diluted to 1 ml with buffer 106 and PEG precipitation buffer in a 1:1 ratio. The mixtures were centrifugated at 17,800 x g for 25 min in room temperature. The pellet was resuspended in buffer 106, and the final concentration was adjusted to 20 nM. For dimerization the solutions were mixed in a 1:1 ratio incubated at 35 °C with slight shaking for at least 14 hours.

### Development of the dimerization procedure

In our initial attempt to connect the bottom and top disks, we used a multi-valent connection that consisted of the swivel hybridization site and 2-4 additional hybridization sites located at the foothold positions. This configuration, however, resulted in higher order byproduct structures (data not shown). To avoid such byproducts, we resorted to connecting the origami disks using the swivel interaction alone, a reaction that has previously resulted in ∼14% yield for a similar rotor construct^10^. Following optimization of the reaction parameters, which included purification to remove truncated swivel strands, purification to remove excess swivels, and optimization of cation and origami concentrations, temperature, and incubation time, the procedure resulted in ∼70% yield (Fig. 2B).

### Synthesis of custom-sized gold nanorods

Nanorods with size 40 nm x 80 nm were synthesized according to a published method^11^ and stored in 3 mM CTAB at room temperature. The nanorods were characterized by their UV-Vis extinction spectra and by transmission electron microscopy (TEM). The size distribution of the nanorods was estimated using Image-J and custom-written analysis software (Matlab).

### Functionalization of gold nanorods with DNA

The gold nanorods were functionalized with thiolated 21T DNA using a salt aging method^12^. For each nanorod solution unit (1 ml, Absorbance∼1.5, 3 mM CTAB), 5 µl of 100 mM TCEP solution was added to 40 µl of 100 µM thiol solution (water suspension) and incubated for at least 30 minutes. Each unit of nanorod solution was centrifugated at 6000xg, and 0.9 mL of the supernatant was replaced with water. The solution was centrifugated again, the supernatant was carefully fully removed, and 45 µL thiolated DNA-TCEP mixture was added to the pellet and incubated for 10 minutes. An equal volume of 2x TAE, 0.1% SDS was added and incubated for 10 minutes, followed by increasing the NaCl concentration to ∼0.7 M over the course of ∼2 hours. First a salt aging buffer (1x TAE, 1 M NaCl, 0.1% SDS) was added with volumes of 5, 5, 5, 7, 7, and 7 µl. Between additions, the solution was vortexed and spun down gently, sonicated for 5 seconds, and incubated for 10 minutes. Two final salt aging steps consisted of 10 µl of 5 M NaCl, with 30 minutes of incubation between steps. The procedure was usually scaled up by pairing units after the nanorod and thiolated DNA mixing step and adjusting subsequent volumes accordingly. After overnight incubation, the solution was centrifugated four times and resuspended in buffer 102. The final concentration was adjusted to ∼10 nM.

### Attachment of nanorods to the rotary motor

The rotor solution was adjusted to 10 nM, mixed with the nanorod solution (10 nM) in a 1:5 ratio, and incubated for at least 20 minutes at room temperature.

### Rotor immobilization in the microfluidic device

First, the glass surface of a working chamber of a microfluidic device was prepared with rotor immobilization components (BSA and Neutravidin), incubated for several minutes, and washed with T50 buffer. The concentrations of BSA and Neutravidin were 1 mg/ml and 0.2 mg/ml, respectively. The solution of rotors coupled to gold nanorods was diluted with 502 buffer to ∼0.2 nM and flown into the operation channel. After the desired concentration was reached, the channel was washed with 102 buffer. To achieve full anchoring of the bottom disks, an additional step of low concentration (100 nM) biotinylated strand addition was performed. These strands hybridized to vacant DNA anchoring sites emanating from the bottom disks and secured their attachment to the Neutravidin-coated surface.

### Agarose gel analysis of origami assemblies and gold nanorod assemblies

The samples were electrophoresed on 1% agarose gels containing 100 mM Na^+^ and 2 mM Mg^2+^ in an ice water bath for 90 minutes at 45V.

### Transmission electron microscopy

For sample preparation, 300-mesh copper grids (Ted Pella, Prod No. 01813-F) were glow discharged, then 2.5 µL of sample was applied to the grid. After 1 minute, excess liquid was removed by blotting with filter paper. The grid was dried in air for 1 minute, followed by application of 5 µL of 2% uranyl acetate (SPI CAS# 6159-44-0) as the negative stain for samples containing origami. The grid was blotted to remove the excess uranyl acetate and was dried in air before insertion into the microscope. The imaging was performed using a Talos F200C TEM microscope (ThermoFisher Scientific) operating at 200 kV equipped with a Ceta 16M CMOS camera.

### Atomic force microscopy imaging

Data were acquired with a Cypher instrument (Asylum Research). Structures were deposited on freshly cleaved mica and imaged in a 1xTAE buffer containing 12 mM MgCl_2_.

### Cryo-electron microscopy measurements

#### Grid preparation

Grids were prepared by plunge-freezing into liquid ethane. Aliquots of 3 µl of sample solution at a concentration of ∼200 nM were deposited on glow-discharged Quantifoil R 1.2/1.3 holey carbon grids (Quantifoil Micro Tools GmbH). Samples were manually blotted for 3 seconds at room temperature and rapidly plunged into liquid ethane using a home-built plunging apparatus. The frozen samples were stored in liquid nitrogen until imaging.

### Data collection

The prepared grids were loaded onto a FEI Tecnai F30 Polara microscope (FEI, Eindhoven) operated at 300 kV, under cryogenic conditions in a low-dose mode. Datasets were collected using SerialEM, using a homemade semi-automated data collection script. Images were collected by a K2 Summit direct electron detector fitted behind an energy filter (Gatan Quantum GIF) set to ±10 eV around zero-loss peak. Calibrated pixel size at the sample plane was 2.3 Å. The detector was operated in a dose fractionated counting mode, at a dose rate of ∼8 ē/pixel/second. Each dose-fractionated movie had 50 frames, with total electron dose of 80 ē/Å2. Data were collected at a defocus range of -1.0 to -2.0 μm.

### Data processing

Data processing was done using Relion3.1. Movies were motion corrected using Relion3.1 motion correction implementation, and estimation of the defocus values was done by using Gctf. Particles were initially picked based on LoG autopicking, followed by a few iterations of template-based 2D classification that removed “bad” particles. The chosen 2D classifications classes were used to generate an initial model, followed by 3D classification to further divide the data into useful groups from which to choose. The chosen 3D classifications classes were later refined to create a final Cryo-EM map.

## Section 6: Rotor Brownian Rotation

### Mobility as a function of cation concentration

In the Brownian rotation state (i.e., in the absence of fuels), some of the observed rotors were immobile. The mobility of the rotors was inversely proportional to the concentration of cations in the working buffer. For example, when repeatedly switching between a buffer with a high cation concentration (1000 mM Na^+^) to a buffer with a low cation concentration (100 mM Na^+^ and 2 mM Mg^2+^), some rotors switched from an immobile to a mobile state (Fig. S4A, purple and grey incubation periods, respectively). Analysis of a set of ∼300 rotors showed that only 40% of the rotors were mobile in the high concentration buffer, whereas 75% were mobile in the low concentration buffer (Fig. S4B, left panel). The root mean square deviation of the rotors, which is an indication of their degree of mobility, also depended on cation concentration (Fig. S4B, right panel). Overall, the buffer with cation concentrations of 100 mM Na^+^ and 2 mM Mg^2+^ resulted in the highest mobility and was selected for rotor operation.

### Dwell time analysis

The mobility of the rotors in the Brownian rotation state in the selected working buffer was also characterized in terms of the dwell times (durations) in temporary, immobile states. A histogram of the maximum dwell times during 2 hours of rotor operation (of a set of ∼150 mobile rotors) showed that for most of the rotors the maximum dwell time was between seconds to 3 minutes; the maximum dwell time was rarely 10 minutes or above (Fig. S4C, D).

**Figure S4.**
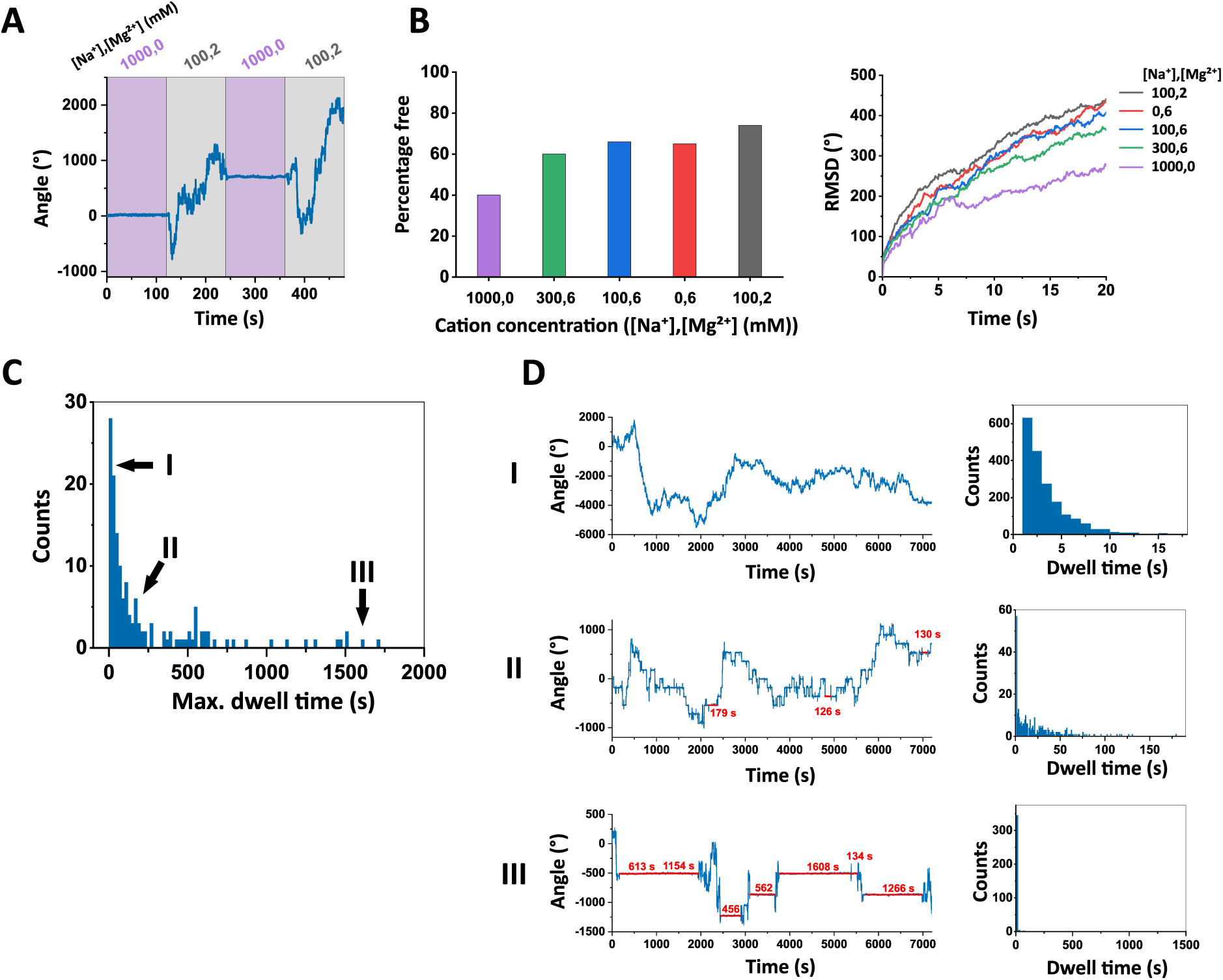
Brownian rotation. **(A)** Time-trace of a rotor in the Brownian rotation state as the working buffers were switched between high (purple) and low (grey) cation concentration buffers. The left and right numbers in the notation above the incubation times refers to Na^+^ and Mg^2+^ concentrations (mM). **(B)** Percentage (left panel) and root mean square deviation (RMSD; right panel) of freely rotating rotors as a function of cation concentration in the working buffer. Rotors were considered freely rotating if the standard deviation of the Gaussian fitted to the time-trace segments of 20 seconds was larger than 10°. **(C)** Histogram of maximum dwell time (i.e., immobile state duration) per rotor in a 2-hour measurement accumulated from 5 fields of view, 2 recordings each (automatic scanning). Segments constrained within 70° were considered dwells. **(D)** Left: Time-traces of three exemplary rotors marked by I, II, and III in panel C with maximum dwell times of 15, 179, and 1608 seconds, respectively. Dwells longer than 120 seconds are marked in red on the trace. Right: Dwell time histograms of the three rotors.

## Section 7: Rotor Operation

### Binding to the track

Starting from the free Brownian rotation state, introduction of a fuel command allows the rotor’s leg to bind to the corresponding foothold through the fuel (Fig. S5A, top route). The antifuel then returns the rotor to the Brownian rotation state. Such operations were successfully performed by the rotor using each of the six fuel and antifuel pairs as shown in Fig. S5B. Fig. S5C shows a rotor subjected to 10 such cycles, successfully binding and unbinding the track nearly 60 times over the course of ∼12 hours. Track binding failure is expected due to the possible scenario of two separate fuels binding the leg and the foothold instead of a single fuel joining them, leaving the rotor in a free rotation state (Fig. S5A, bottom route).

### Rotor initialization with all legs bound to the track

While initialization at state [1,2] is possible through the top route in Fig. S5A, to avoid unsuccessful leg binding reactions due to double fuel binding, the initialization of the rotor in state [1,2] for directional rotation was typically performed using an alternative mechanism involving leg blocking (Fig. S6A). First F6 is inserted to block L2, followed by F2 and then AF6 to release the leg, allowing it to bind to T2. Next a similar procedure is used for binding T1 using F3 as a blocker. Fig. S6B shows a typical response to the procedure. The rotor does not behave exactly as shown in Fig. S6A due to the inherent stochasticity, but its arrival at state [1,2] is evident.

An alternative method to avoid double fuel binding with “direct fuel addition” as in Fig. S5A is to decrease the concentration of fuel, thereby decreasing the fuel binding rate relative to the leg placing rate. It is also possible to skip the initialization step completely and apply the FBAF rotation sequence directly from the Brownian rotation state. By starting, for example, with F3, AF1, F4, AF2, etc., the rotor is expected to be fully initialized at state [4,5] after double fuel binding event possibilities are used up.

**Figure S5.**
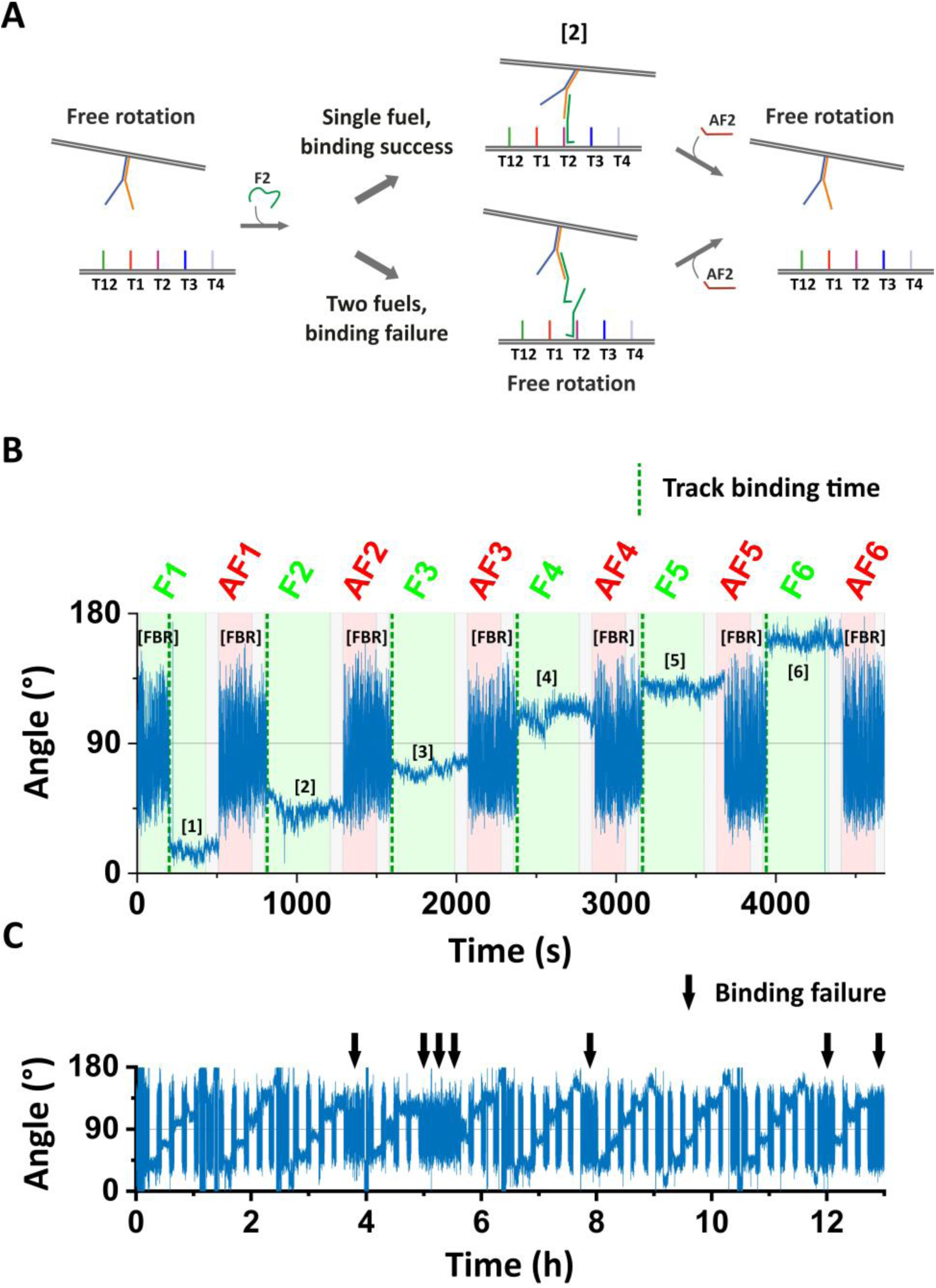
Track binding reactions. **(A)** Schematic of the track binding and unbinding operations including a successful track binding path (top) and a failed reaction path due to double fuel binding (bottom). **(B)** Exemplary rotor time-trace of six successful track binding reactions. The state of the rotor after each reaction is denoted in black in the time trace. Track binding times are denoted by dotted green lines and were extracted by an automatic script. **(C)** Ten cycles of the experiment performed in panel B, 60 reactions in total. The black arrows denote failed track binding reactions in which the rotor remained in the Brownian rotation state.

**Figure S6.**
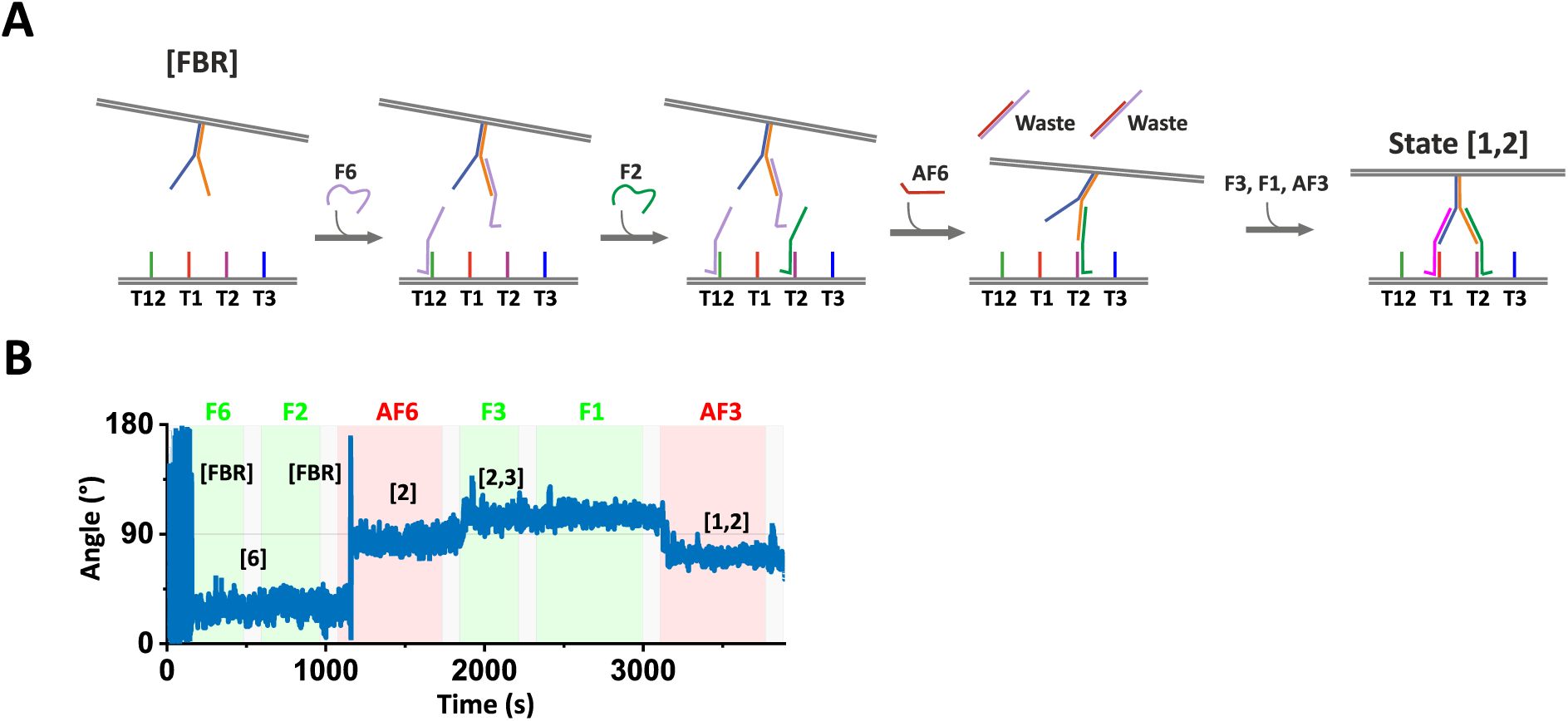
Rotor initialization. **(A)** Procedure to transfer the rotor from a free Brownian rotation state to state [1,2] using blocking to avoid double fuel binding of the destination state fuels. **(B)** Typical response to the procedure.

**Figure S7.**
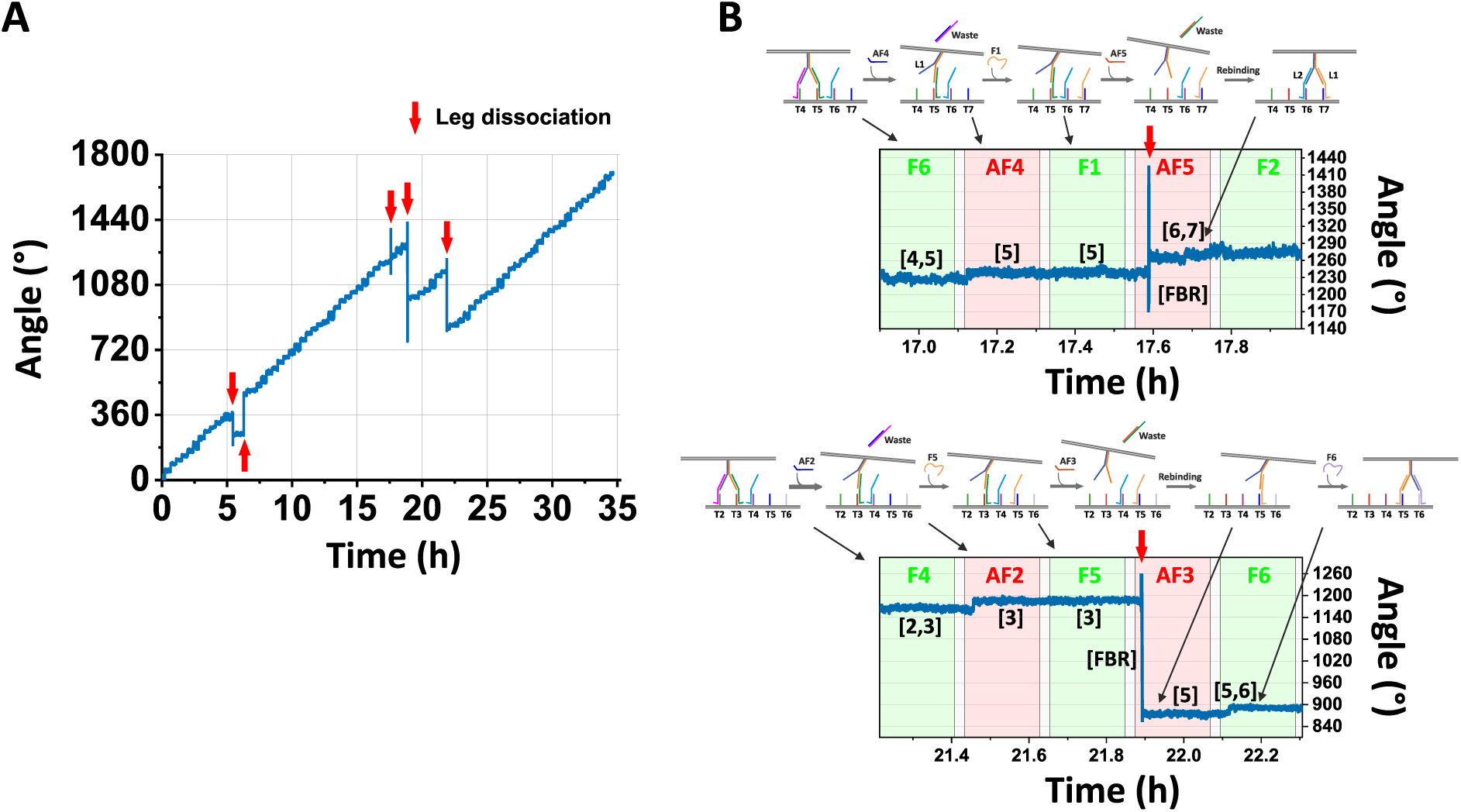
Rotor errors and recoveries. **(A)** Exemplary rotor time-trace showing clear dissociations from the track (denoted by red arrows), in which the rotor rebound to the track in phases with a size of a multiple of half of a rotation in accordance with the symmetry of the circular track. **(B)** Additional examples of dissociations from the track with mechanistic details for the rotor in analyzed in panel A. The errors selected are indicated by the time axes. The durations of the free Brownian rotation states until rebinding were 5 and 10 seconds for the top and bottom traces, respectively.

### Stepping reaction profile fitting

The stepping reaction profiles were fitted with two simple kinetic models. The first is a double exponential model that describes a combination of fast and slow rotor steps, detailed in the following equations:

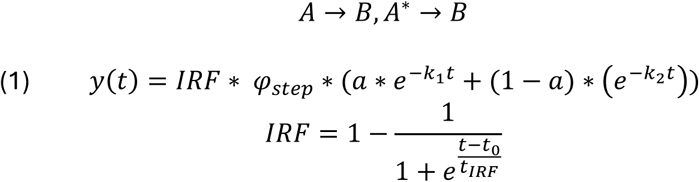

where 𝐴 and 𝐴^∗^ represent two distinct rotor populations in the origin state; 𝐵 is the destination state; 𝜑_𝑠𝑡𝑒𝑝_ is the average step size; 𝑎 is the fraction of population; 𝐴, 𝑘_1_, and 𝑘_2_ are first order rate constants; t is the time from insertion of the antifuel command; IRF is the instrument response function; 𝑡_0_ is the time of arrival of the antifuel command wavefront to the observed position in the microfluidic working chamber; 𝑡_𝐼𝑅𝐹_ is the instrument response time (indicating the speed of solution replacement once the wavefront has arrived); and 𝑦(𝑡) is the average angular progression.

The second model is the consecutive reactions model, which accounts for separate leg lifting and leg placing operations, as detailed in the following equations:

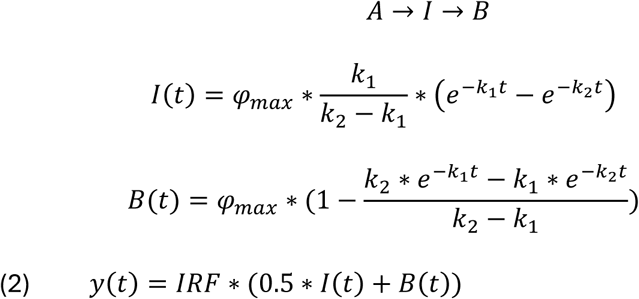

where 𝐴 is the origin state; I is an intermediate state; 𝐵 is the destination state; 𝜑_𝑠𝑡𝑒𝑝_ is the average step size; 𝑘_1_and 𝑘_2_ are first order rate constants; and 𝑦(𝑡) is the average angular progression, in which y considers the contribution of intermediate state I to the angular progression.

Neither calculated curves fit the data well. In the first 35 seconds, the data is in several cases faster than the fitted curves (Fig. S8). This, however, is reasonable, as the behavior of the rotary motor is influenced by more factors than those considered in the models such as uncorrelated origami body movements and walker operations due to origami body lagging, and separate response times of the two bipedal walkers. More accurate fitting will require integrating these factors into the kinetic models.

**Figure S8.**
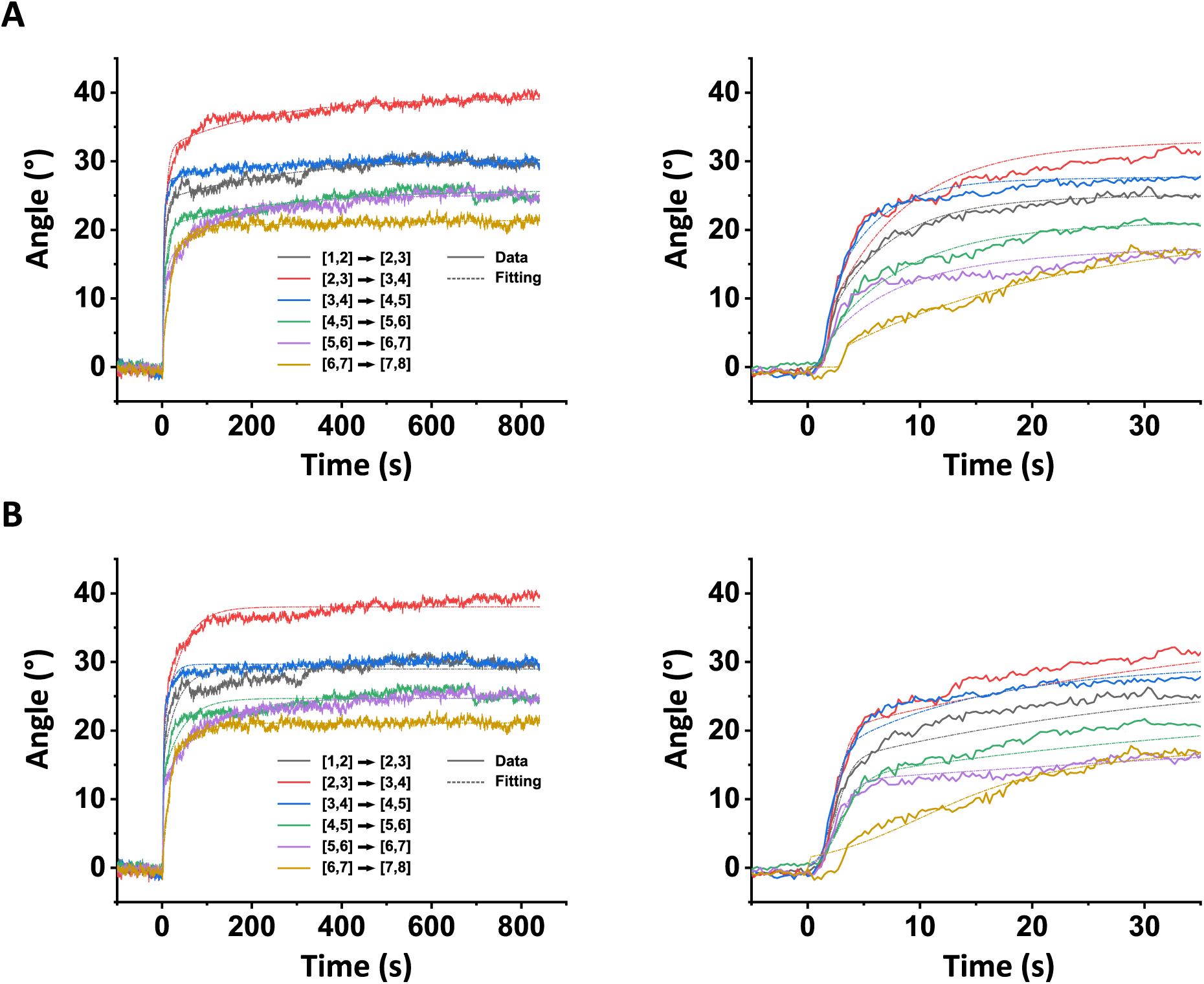
Stepping reaction profile curve fitting. **(A)** Double exponent model. **(B)** Consecutive reactions model. Left panels: full step time-traces. Right panels: First 35 seconds.

**Figure S9.**
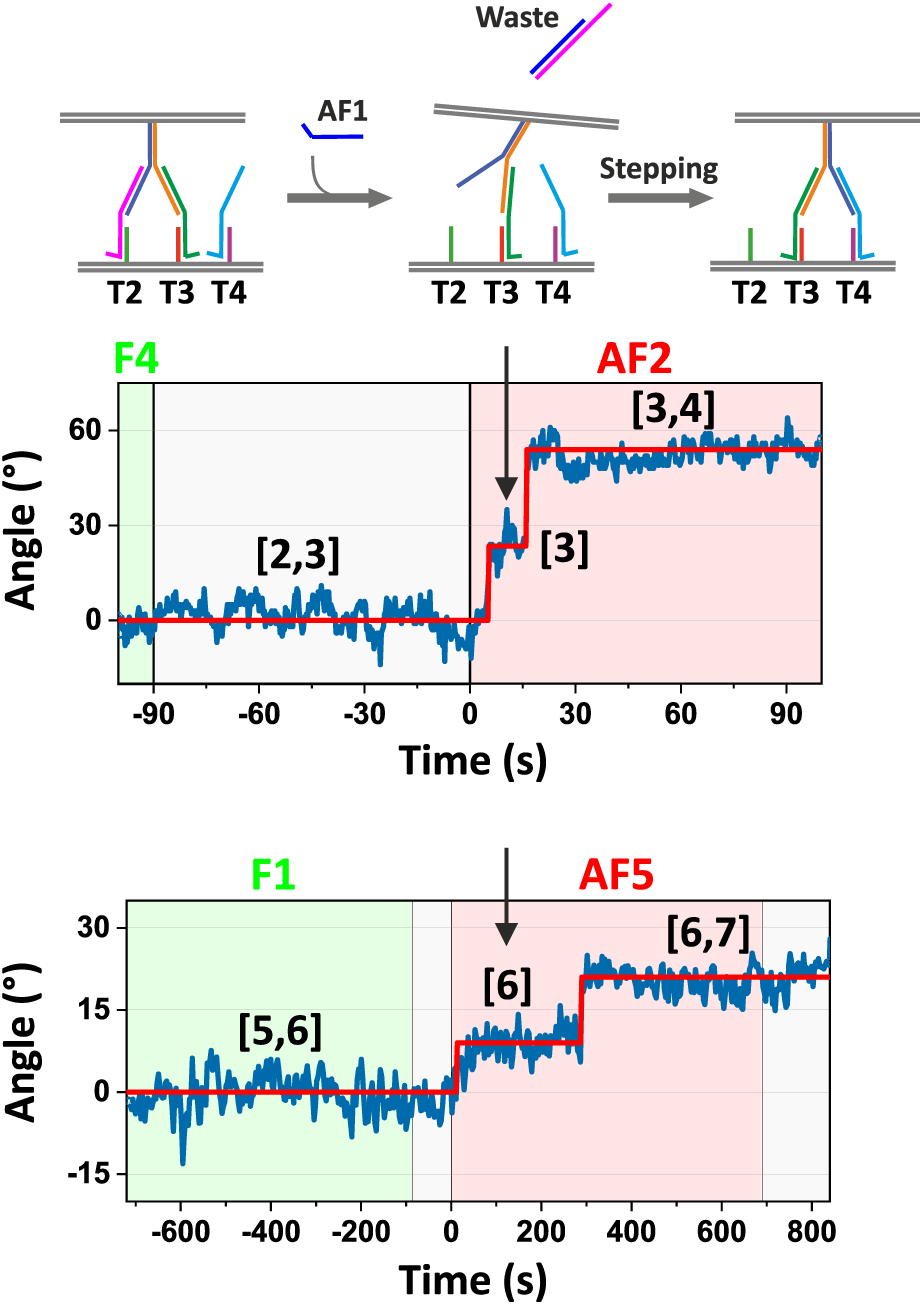
Step fitting. Shown are time-traces of two exemplary rotor steps with fitted traces in red. The function used for the fitting was a step function with a possible intermediate state. Steps with fitted total sizes in the range of 10°-70° and with fitted responses within 0-650 seconds of the antifuel command insertion time were used for further processing. In both of the cases shown here, an intermediate was detected. In the second case, an intermediate state separated by as little as 10° from the original state was clearly detected. A schematic of an interpretation of the time-trace where the intermediate state is attributed to a leg-lifted state is shown on top. Rotor states during the reaction are given in black text over the steps.

### Differences in rates between fuels

The normalized kinetic profiles of the stepping reactions show large rate differences between the six different steps of up to 10 fold (Fig. S10A). Within a few seconds, 70% of the [3,4]→[4,5] steps were complete, whereas only after ∼30 and ∼60 minutes were the same percentages of steps completed for the [6,7]→[7,8] and the [5,6]→[6,7] steps, respectively.

Differences between fuels also occurred in the track binding reactions in which freely rotating rotors respond to fuel commands by binding to the track (Fig. S10B). For most fuels the responses occurred within 10-30 seconds, but for F1 the responses were much slower and required up to hundreds of seconds.

Interestingly, both the F1 track binding reaction and the [5,6]→[6,7] stepping reaction involve the same leg placing reaction on T1 and T7, which suggests that either the leg placing reaction is slow or the rotor has difficulty accessing the orientations of these footholds. To test the effect of rotor orientations on the efficiency of the track binding reactions, we performed the track binding reactions with several rotor variants (Fig. S10C): the original variant containing all walkers and footholds, the four variants containing a single set of six footholds and a single walker, a variant with a walker in a position with a 90° shift (in the position of T12 instead of T3\T9), and a variant where two footholds were inverted in position (T2 and T5). In all cases, the reactions resulted in approximately the same fuel binding efficiency curve (Fig. S10D), indicating that the reason for differences between fuels in kinetics is not the positions where the reactions occur but rather the sequences of the machinery involved in the reactions.

**Figure S10.**
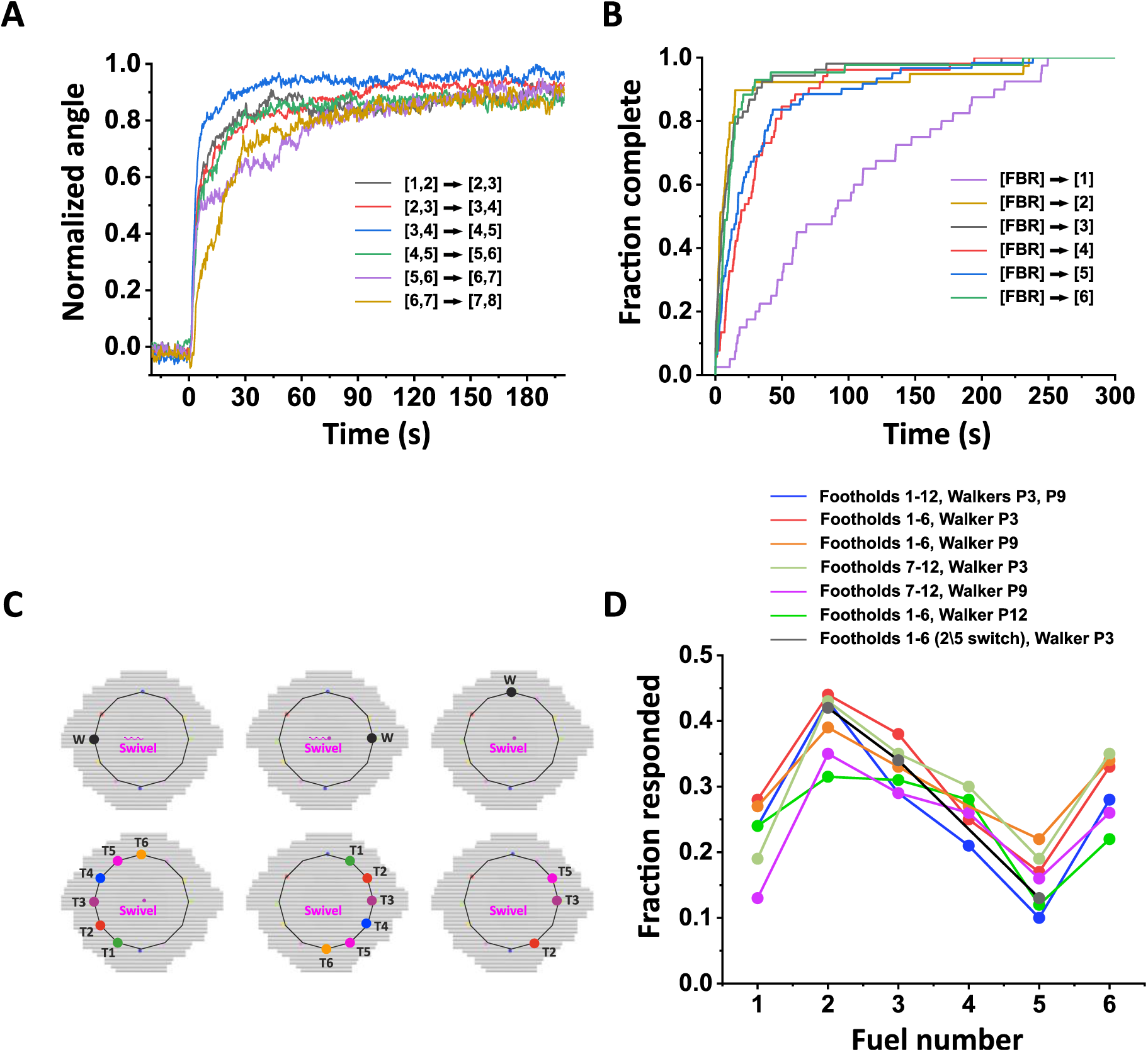
Differences in rates between fuels. **(A)** Angles as a function of time for ∼120 steps for each of the six step types as in Fig. 5A, normalized with the maximum angles obtained from the double exponent model fitting as in Fig. S8A. **(B)** Track binding reaction profiles accumulated from analysis of rotor response times as shown in green dotted lines in Fig. S5B. **(C)** Bottom and top disk variants (bottom and top rows, respectively) used for the track binding efficiency measurements in shown in panel D. **(D)** Track binding reaction efficiencies for rotor variants. Plotted is the fraction of observed rotors (out of ∼300) that responded to indicated fuel within 20 minutes of fuel command introduction (i.e., those that switched from the free rotation state to a track bound state). In this set of experiments the concentration of fuels was 100 nM (in all other measurements the concentration of fuels and antifuels was 10 µM), and for each variant, data from ∼20 fields of view were collected automatically between each of the fuel and anti-fuel commands. Movie segments were 20 seconds long, and rotors were determined to be immobile if the standard deviation of the Gaussian fit to the data was less than 10°.

**Figure S11.**
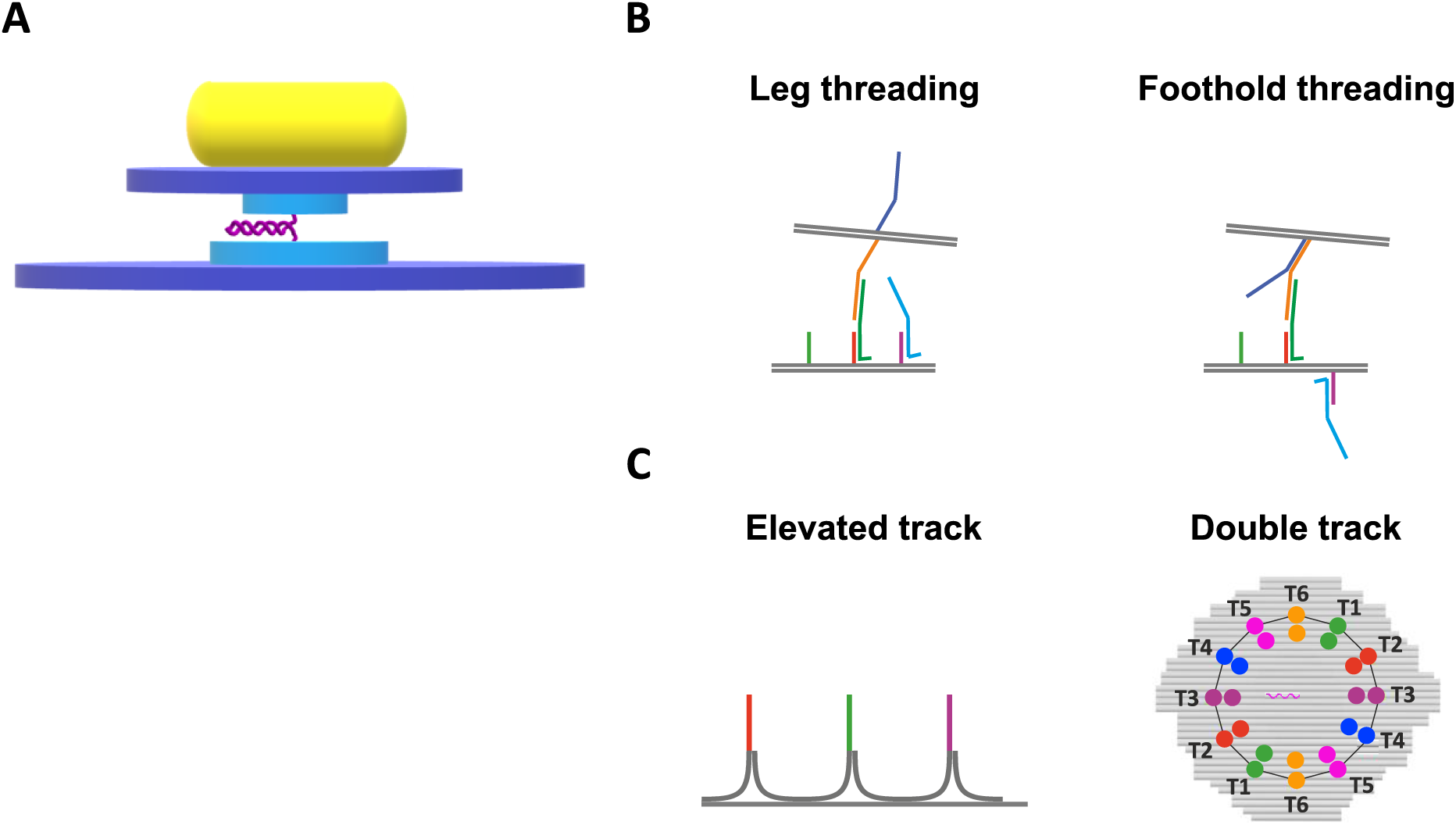
Possible rotor improvements. **(A)** Rotor modified design that should reduce interactions with the surface environment. **(B)** Molecular threading in the rotary motor. **(C)** Design modifications that should counter the threading effect.

**Figure S12.**
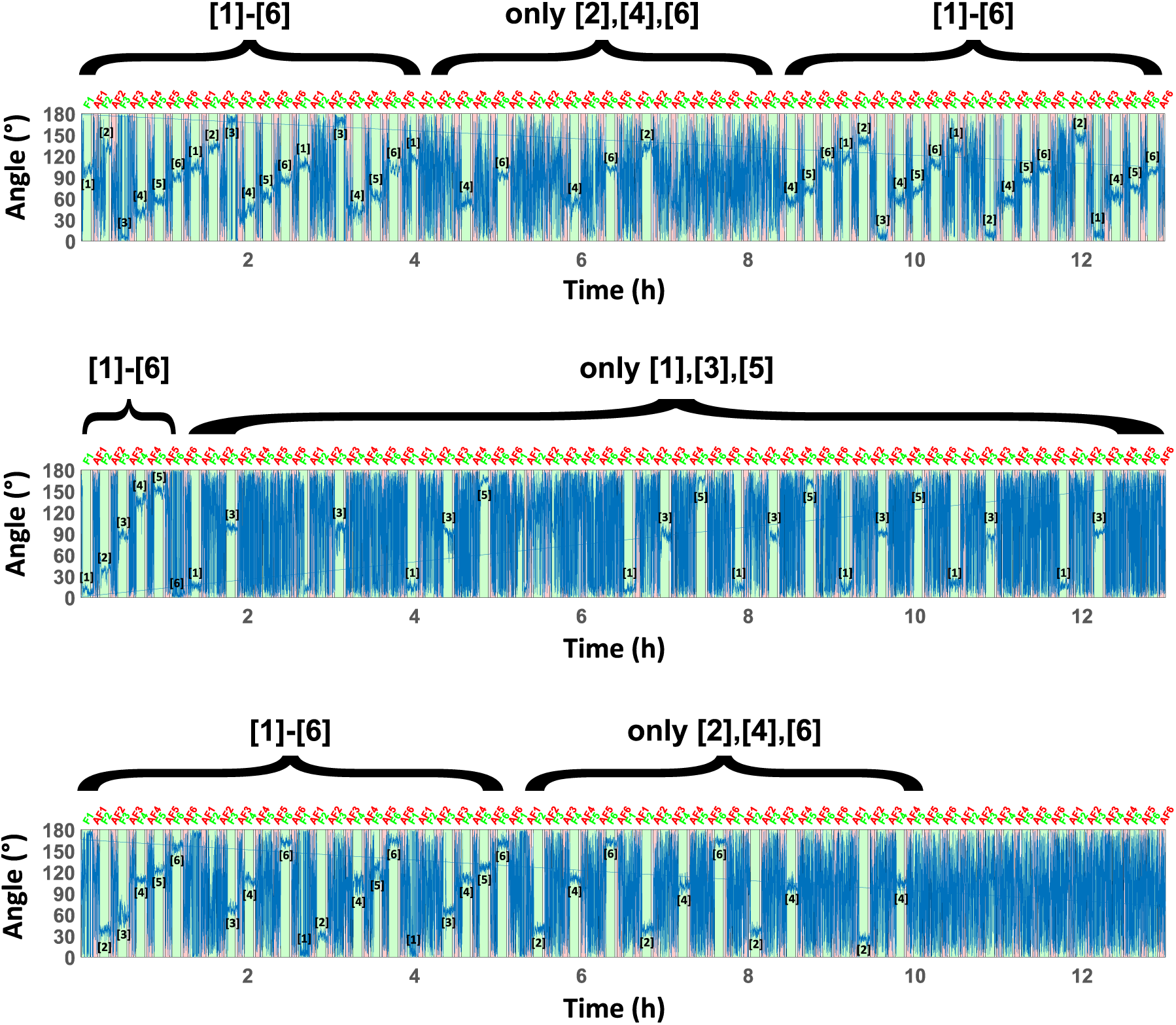
Inactivity of legs. Shown are three exemplary time traces of rotors that failed track binding reactions as denoted in Fig. S5C. Occurrences in which the rotors successfully bound to the track are marked by the corresponding rotor state in black. These rotors were fully active with responses to all 6 fuels in some periods but were only partially active for several hours and had reaction cycles with responses to subsets of even-only- or odd-only-numbered fuels. Such behavior can only be explained by inactivity of legs (which in turn can be explained by interaction with or threading into the origami), as foothold inactivity or difficulty accessing specific orientations is highly unlikely in such scenarios. In the top example the inactivity is temporary, highlighting the reversibility of the interactions.

## Section 8: Supplementary Movies

In all movies, the time-traces contain unbinned data (as opposed to the time traces in the main text corresponding to those shown in Movies 2 and 6).

**Supplementary Movie 1. Rotor operation mechanism.** The rotor’s walkers consume fuels and antifuels to propel the rotor around the circular track.

**Supplementary Movie 2. Single clockwise revolution.** Shown is the rotor from Fig. 3A. For each step, 30 frames are shown.

**Supplementary Movies 3-5. Multiple clockwise revolutions.** The movies correspond to the three time-traces shown in Fig. 3B. For each step, 6 frames are shown.

**Supplementary Movie 6. Rotation in alternating directions.** Shown is the rotor from Fig. 3D. For each step, 6 frames are shown.

## Section 9: Microfluidics Command Sequences

The microfluidics command sequences used for rotor operation are detailed in the following tables:

**Supplementary Table 1.**
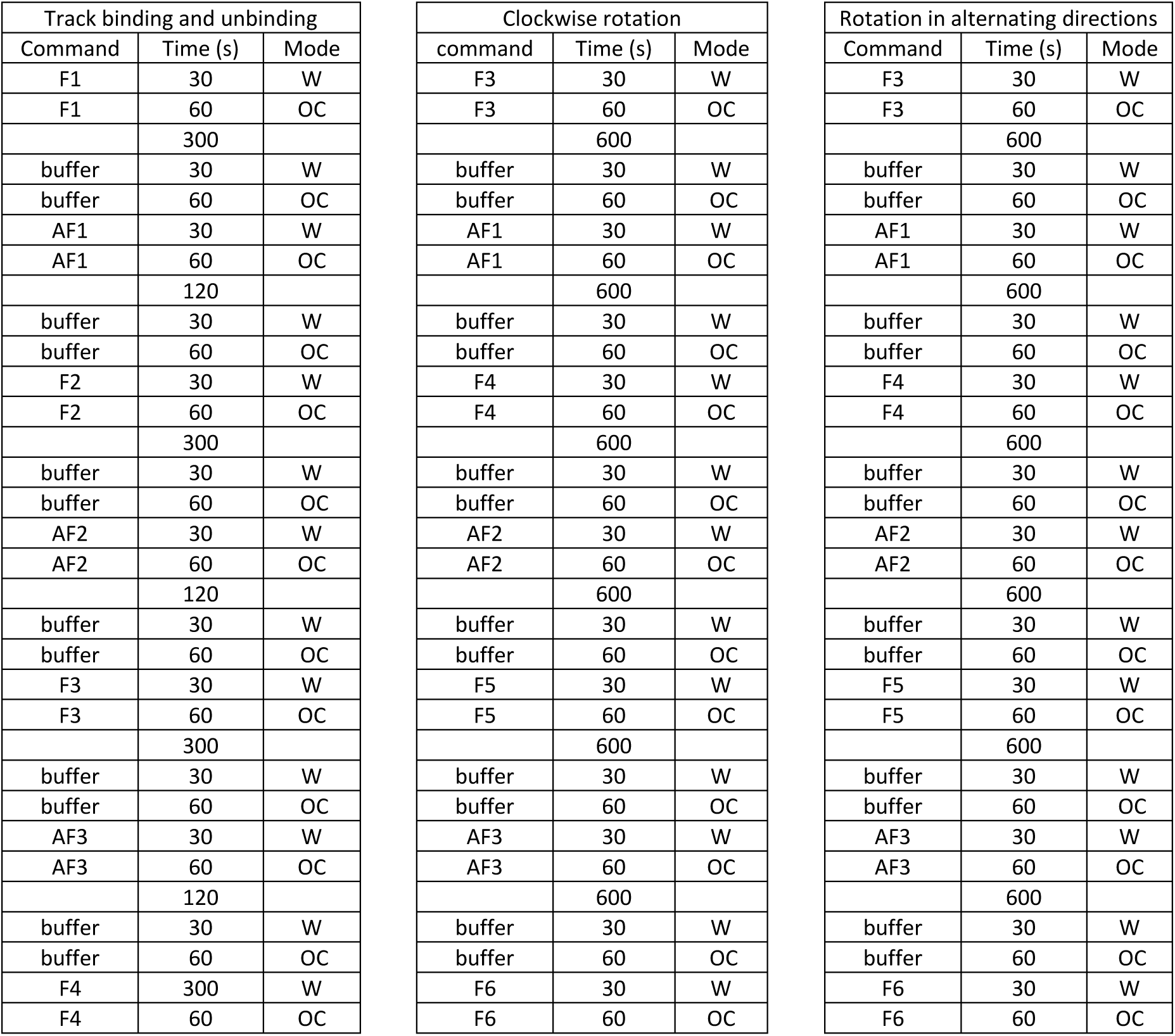
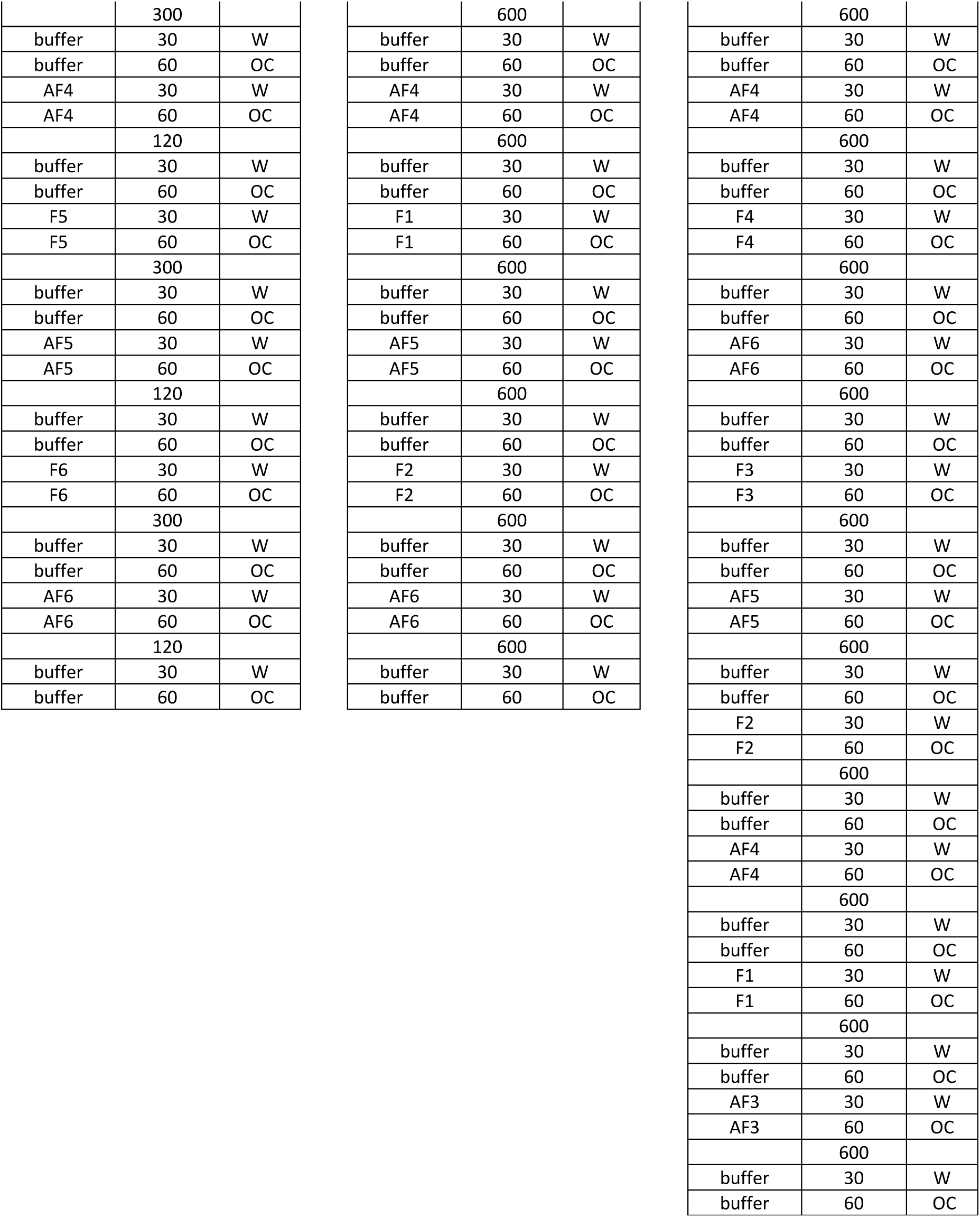
Microfluidics command sequences. Left: Track binding and unbinding (Fig. S5B). Middle: Clockwise rotation (Fig. 3A, B). Right: Alternating directions (Fig. 3D). Operation modes include flow through waste output (W), flow to operating channel (OC), and static incubation (empty cell).

## Section 9: DNA Sequences

### P8064 scaffold

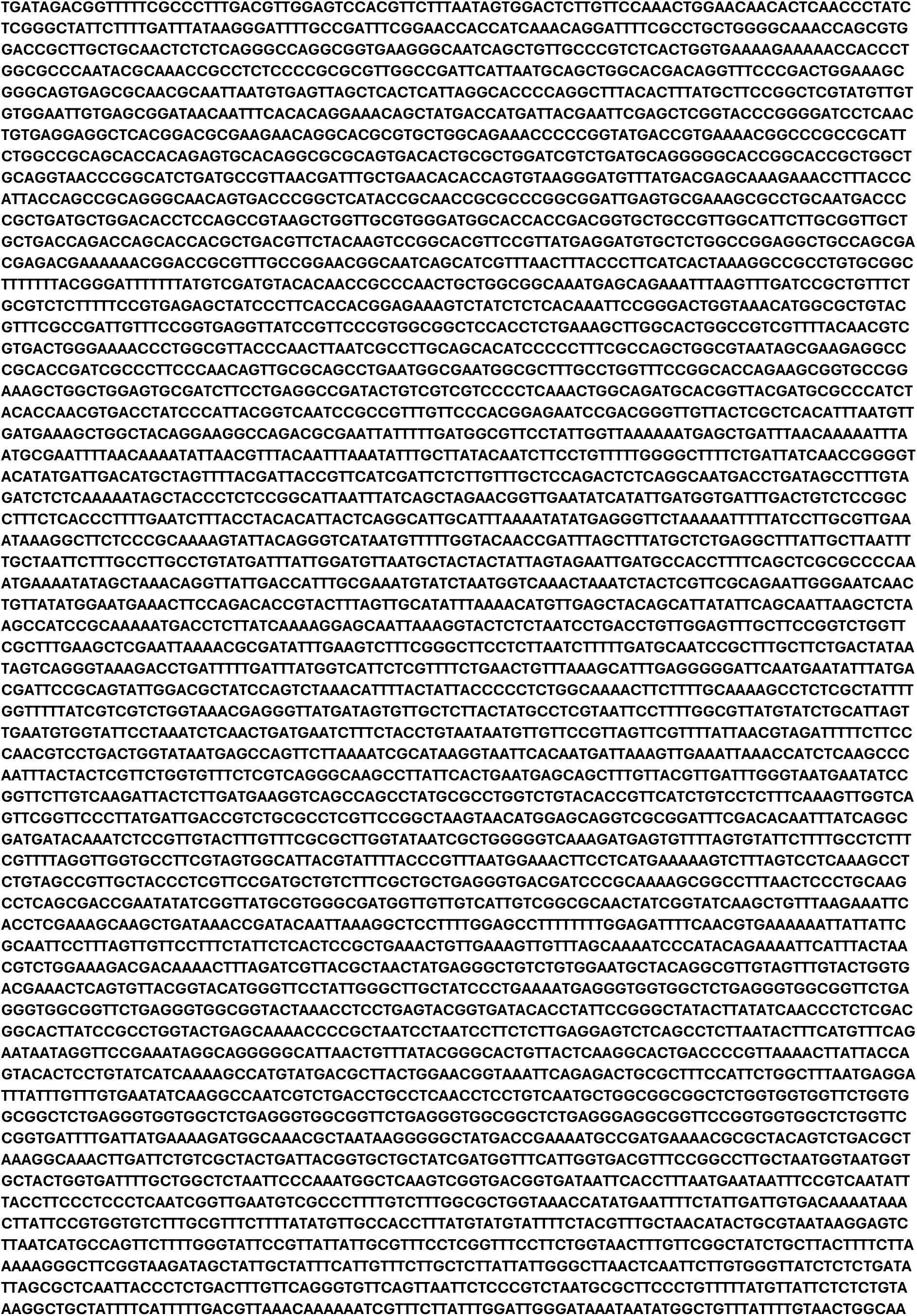
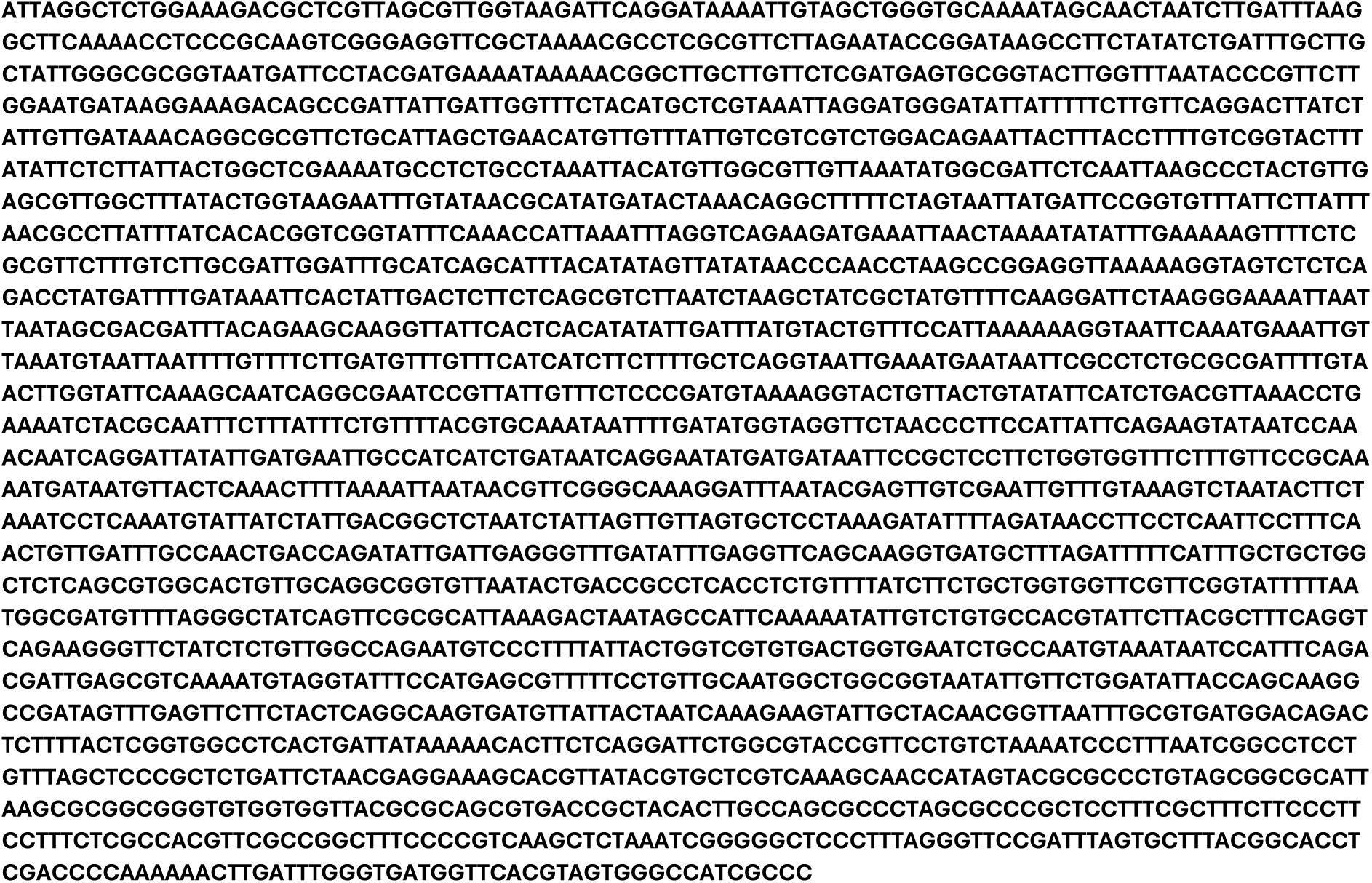

### CS4 scaffold

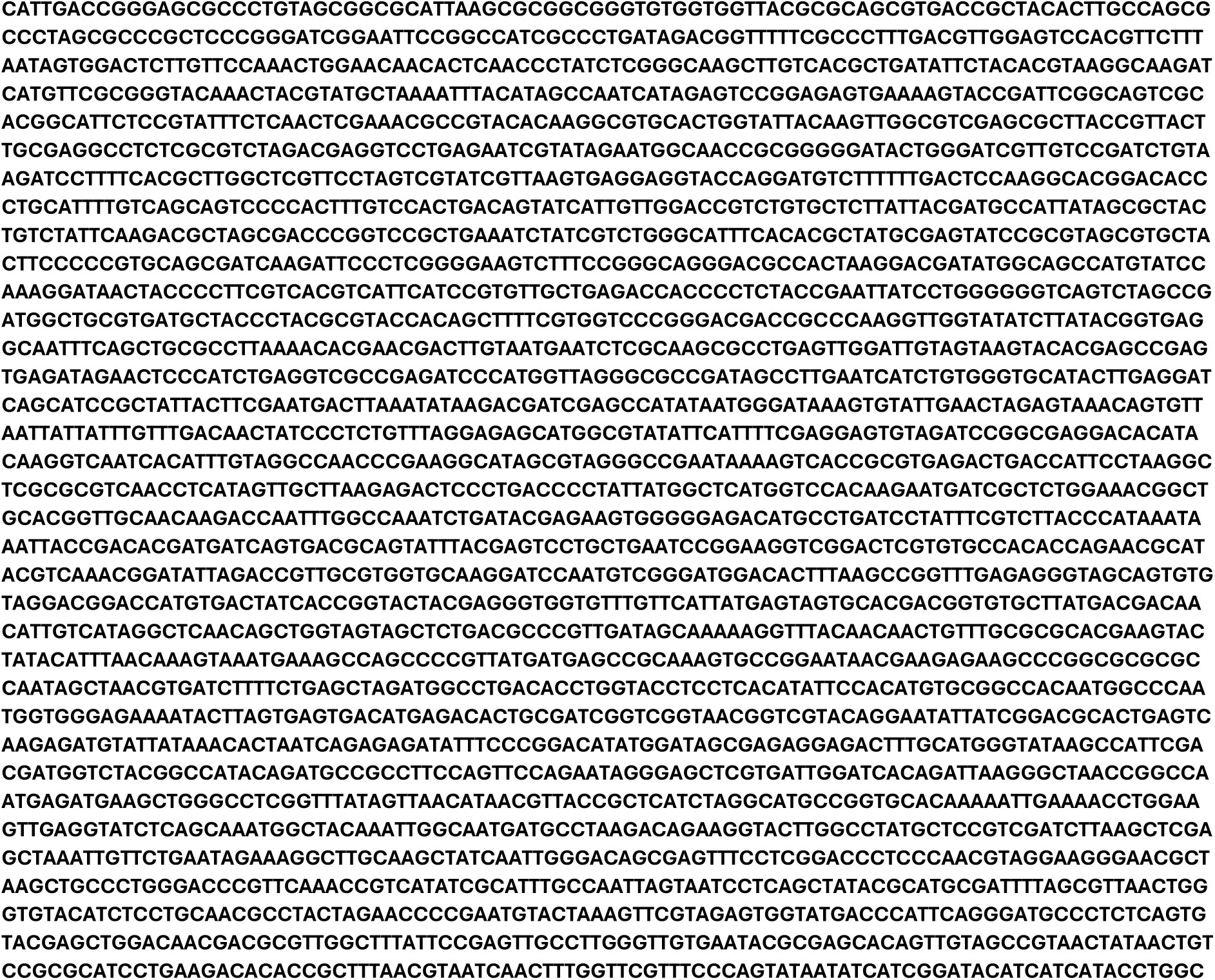
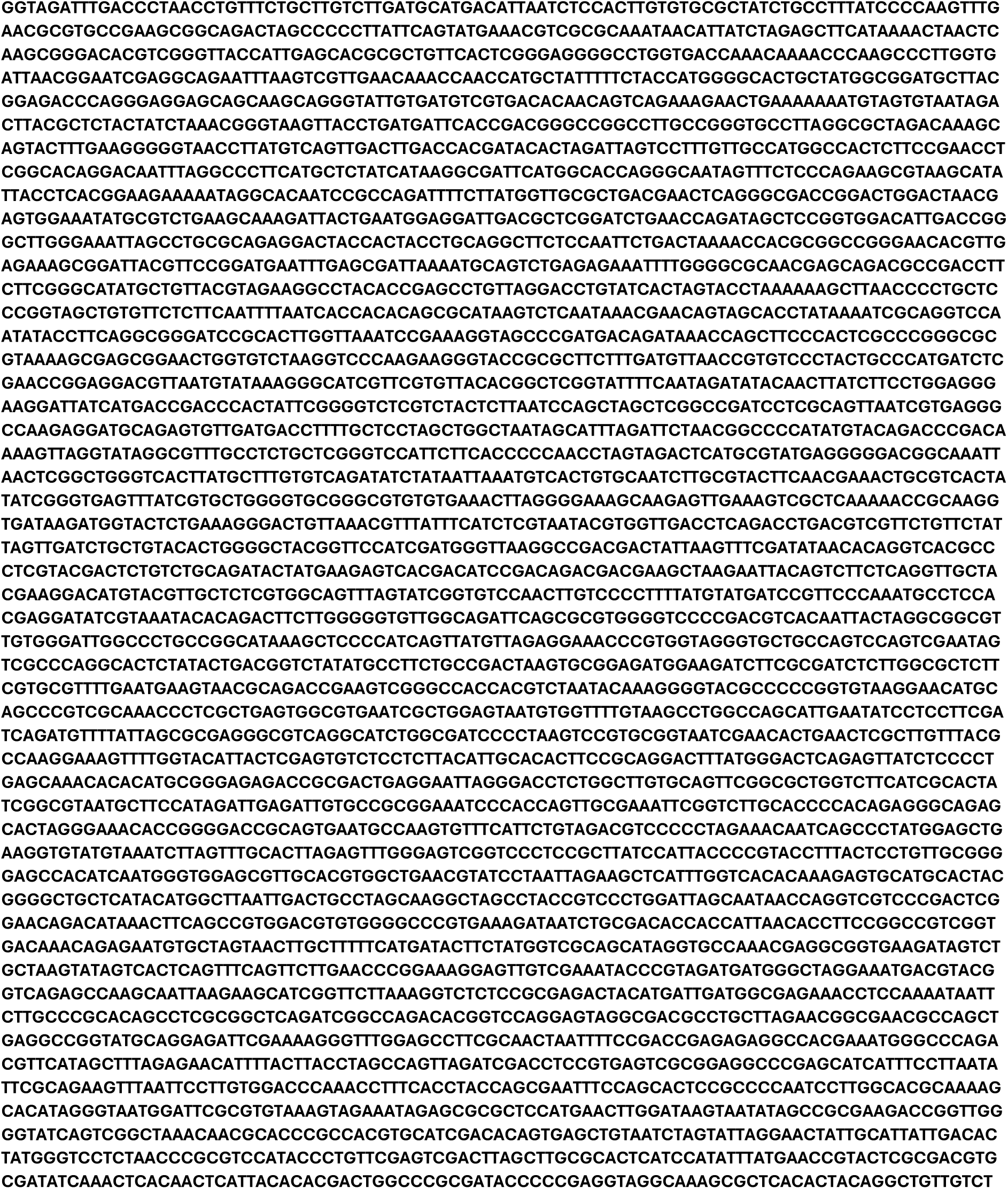

### Staples and special strands

**Supplementary Table 2.**
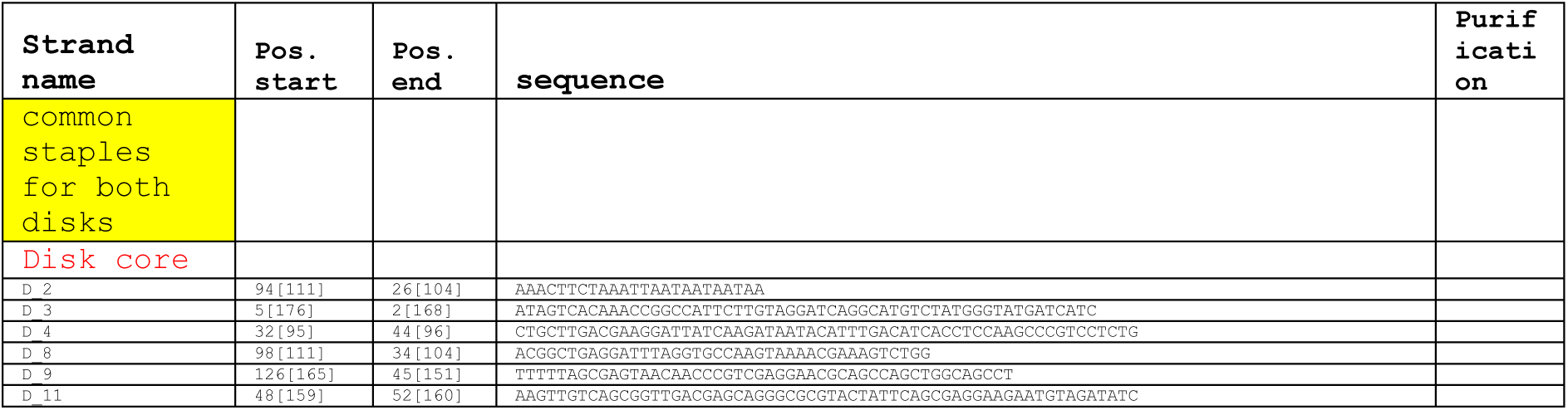
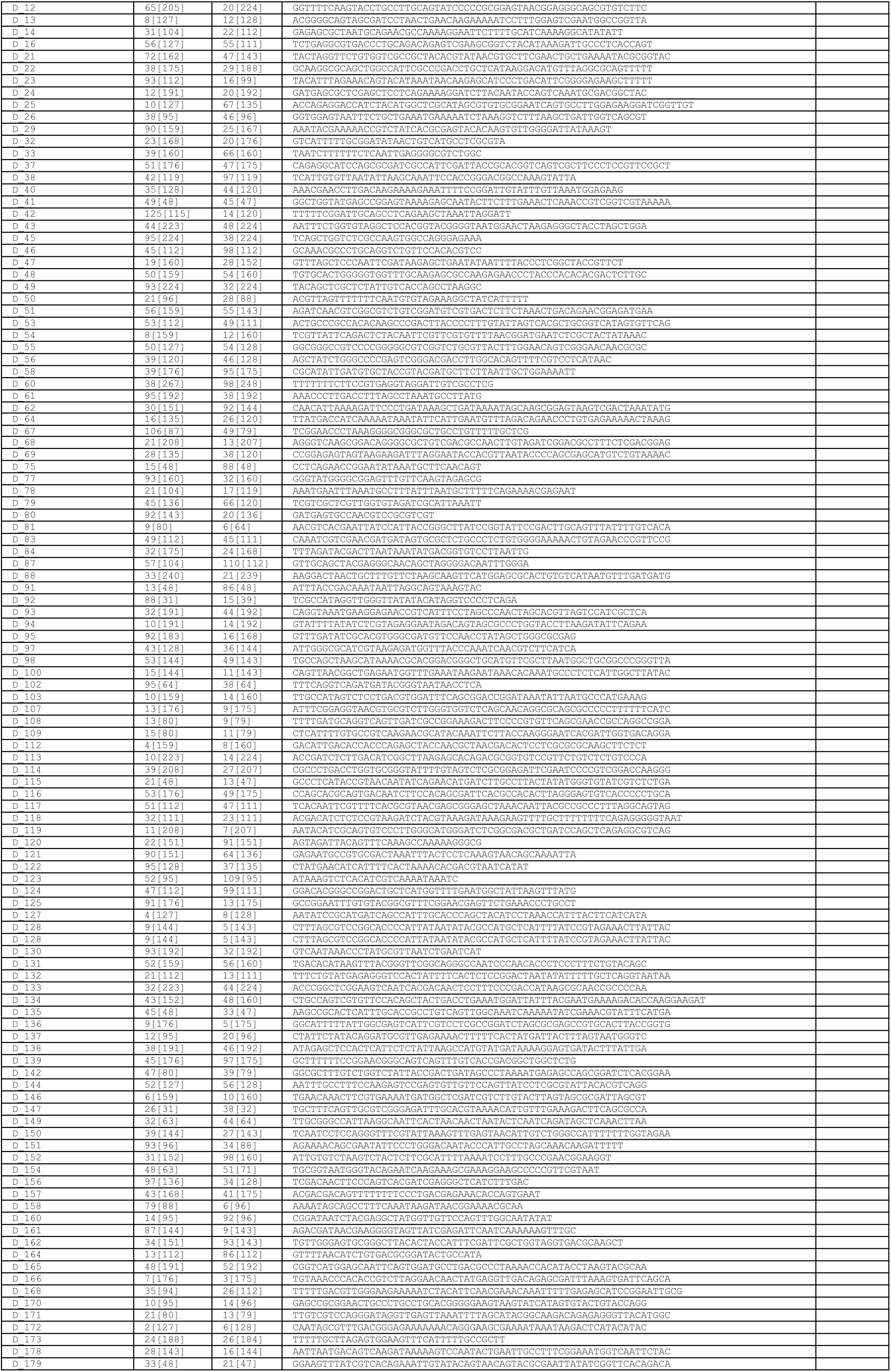
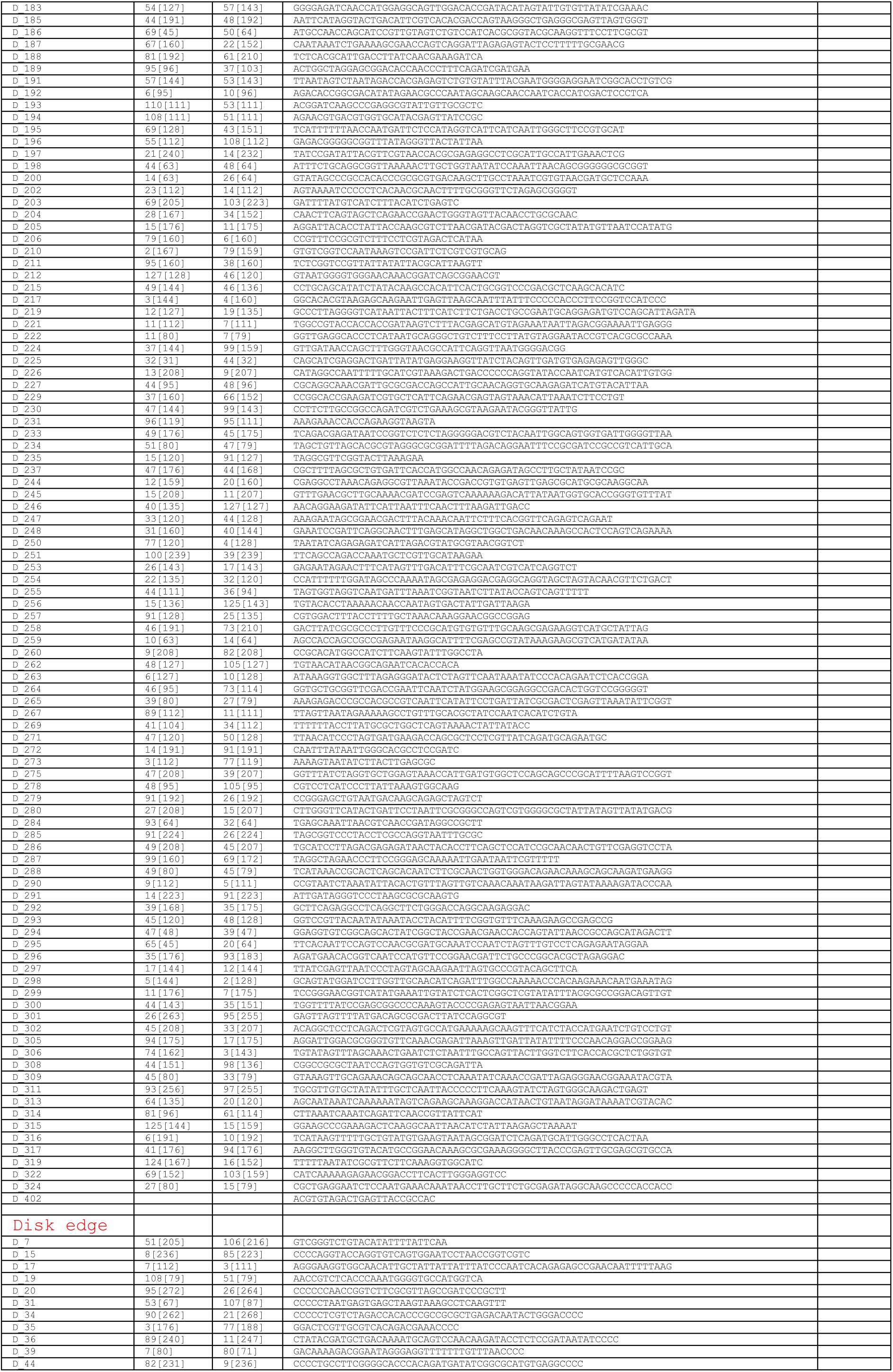
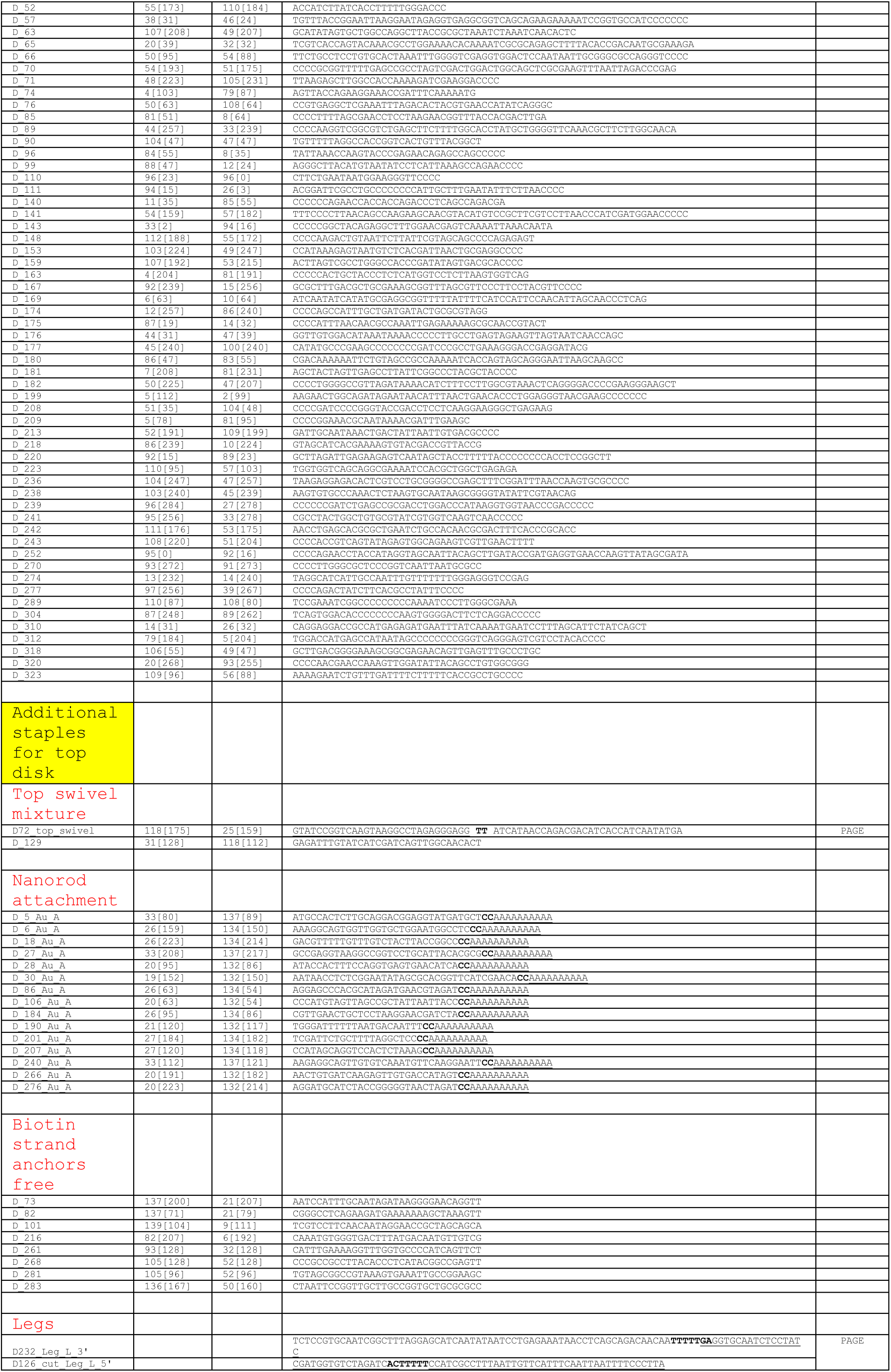
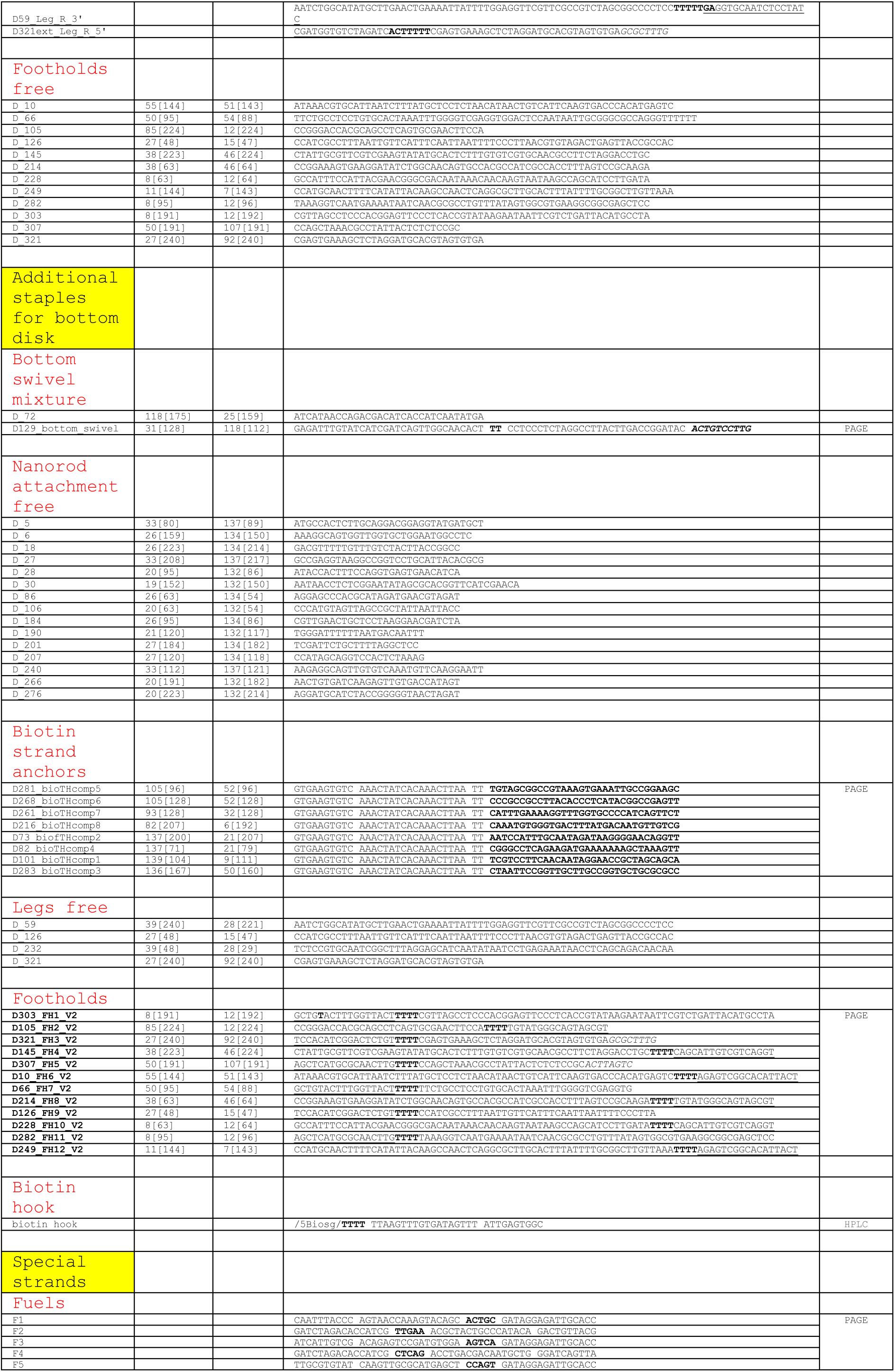
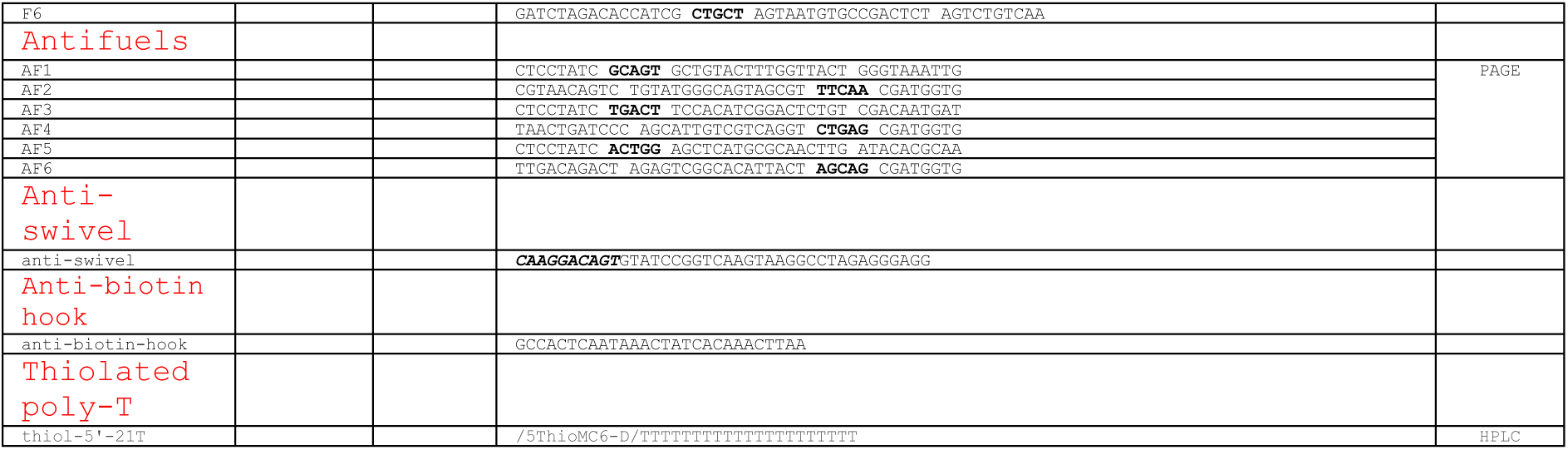
DNA sequences. Disk origami staple mixtures are divided into core staples (used for both disks), top disk special strands and bottom disk special strands. Additional strands used for rotor operation are detailed in the final part.

